# Distinct nucleosome distribution patterns in two structurally and functionally differentiated nuclei of a unicellular eukaryote

**DOI:** 10.1101/018754

**Authors:** Jie Xiong, Shan Gao, Wen Dui, Wentao Yang, Xiao Chen, Sean D. Taverna, Ronald E. Pearlman, Wendy Ashlock, Wei Miao, Yifan Liu

**Affiliations:** Department of Pathology, University of Michigan, Ann Arbor, Michigan 48109, USA; Key Laboratory of Aquatic Biodiversity and Conservation, Institute of Hydrobiology, Chinese Academy of Sciences, Wuhan 430072, China; Institute of Evolution & Marine Biodiversity, Ocean University of China, Qingdao 266003, China; Department of Pharmacology and Molecular Sciences and The Center for Epigenetics, Johns Hopkins University School of Medicine, Baltimore, Maryland 21205, USA; Department of Biology, York University, Toronto, Ontario M3J 1P3, Canada

**Keywords:** Nucleosome positioning, nucleosome occupancy, cis-determinants, trans-determinants, *Tetrahymena*

## Abstract

The ciliate protozoan *Tetrahymena thermophila* contains two types of structurally and functionally differentiated nuclei: the transcriptionally active somatic macronucleus (MAC) and the transcriptionally silent germ-line micronucleus (MIC). Here we demonstrate that MAC features well-positioned nucleosomes downstream of transcription start sites (TSS) likely connected with promoter proximal pausing of RNA polymerase II, as well as in exonic regions flanking both the 5’ and 3’ splice sites. In contrast, nucleosomes in MIC are more delocalized. Nucleosome occupancy in MAC and MIC are nonetheless highly correlated with each other and with predictions based upon DNA sequence features. Arrays of well-positioned nucleosomes are often correlated with GC content oscillations, suggesting significant contributions from cis-determinants. We propose that cis- and trans-determinants may coordinately accommodate some well-positioned nucleosomes with important functions, driven by a process in which positioned nucleosomes shape the mutational landscape of associated DNA sequences, while the DNA sequences in turn reinforce nucleosome positioning.

---

In eukaryotic cells, nuclear DNA is packaged into chromatin, of which the nucleosome is the basic unit, formed by 147 bp of DNA wrapped around a protein core of histones (Kornberg and Lorch 1999). In addition to genetic information carried in DNA, epigenetic information is carried by chromatin, in combinations of histone variants and post-translational modifications (Allis et al. 2007). In addition, nucleosome-associated DNA, in contrast to the linker DNA, may have either decreased or increased access to protein factors, which in various forms bind to DNA, or catalyze reactions using DNA as the substrate or template (Hughes and Rando 2014). Accessibility of genetic information is affected by genome-wide nucleosome distribution patterns, which can be quantified by nucleosome occupancy and nucleosome positioning. Nucleosome occupancy represents the probability of a DNA sequence being associated with any nucleosome in a population, while nucleosome positioning, particularly translational positioning, focuses on the exact 147 bp of genomic DNA occupied by a subset of nucleosomes (Albert et al. 2007; Mavrich et al. 2008). Micrococcal nuclease (MNase) digestion is a well-established procedure for probing nucleosome distribution, based upon its preference to cut the linker DNA, releasing the mononucleosome as an intermediate product (Kornberg and Lorch 1999). MNase-Seq, coupling the partial chromatin digestion with deep-sequencing technologies (Mavrich et al. 2008; Zhang and Pugh 2011), has generated a wealth of information about genome-wide nucleosome distribution patterns in many eukaryotic organisms, revealing various determinants of nucleosome occupancy and nucleosome positioning (Struhl and Segal 2013).

It has long been known that the nucleosome has intrinsic biases for certain DNA sequences (Simpson and Stafford 1983). Poly (dA:dT) tracts are anti-nucleosomal due to their poor bendability (Drew and Travers 1985; Nelson et al. 1987; Segal and Widom 2009), and are generally enriched in nucleosome depleted regions (NDR) (Kaplan et al. 2009; Zhang et al. 2009). Furthermore, strong correlations between nucleosome occupancy and GC-content have been found in in vivo nucleosome distribution of yeast and worm, as well as in vitro nucleosome reconstitution (Tillo and Hughes 2009). On the other hand, sequences with ∽10 bp phasing are often strongly associated with the nucleosome, as the periodic bending facilitates wrapping around the protein core (Satchwell et al. 1986; Kaplan et al. 2009; Zhang et al. 2009; Gaffney et al. 2012). High-resolution mapping of genome-wide nucleosome distribution reveals clusters of nucleosomes, with their dyads offset by ∽10 bp, demonstrating the conservation of rotational positioning (Brogaard et al. 2012).

Other than DNA sequence features, often referred to as cis-determinants, there are also various trans-determinants affecting nucleosome distribution. Transcription in particular has a pervasive influence on chromatin structure, entailing at least partial nucleosome disassembly, even nucleosome eviction at high transcription rates (Weiner et al. 2010). Nucleosome distribution, in energetically favored as well as disfavored DNA sequences, can be further driven by ATP-dependent chromatin remodelers, which actively slide or evict nucleosomes, or exchange histones (Narlikar et al. 2013). All this shapes the nucleosome distribution in gene bodies bounded by transcription start sites (TSS) and transcription end sites (TES), which often manifests as arrays of well-positioned nucleosomes (referred to as +1, +2, etc., ordered 5’ to 3’) (Yuan et al. 2005; Lee et al. 2007; Lantermann et al. 2010; Yen et al. 2012; Hughes and Rando 2014). Importantly, large distances between TSS and the +1 nucleosome strongly suggest promoter-proximal pausing of Pol II (Mavrich et al. 2008; Chang et al. 2012), with emerging significance in transcription regulation (Adelman and Lis 2012).

The relative contributions of cis- and trans-determinants to nucleosome occupancy and nucleosome positioning are actively studied in various eukaryotic organisms (Hughes and Rando 2014; Lieleg et al. 2014). Here we address the effects of transcription in the ciliate protozoan *Tetrahymena thermophila*, taking advantage of its nuclear dimorphism—two types of structurally and functionally differentiated nuclei: the polyploid somatic macronucleus (MAC) and the diploid germ line micronucleus (MIC), in the same cytoplasmic compartment (Karrer 2012). MAC are transcriptionally active, containing largely decondensed chromatin, as well as abundant RNA polymerases and transcription factors (Stargell and Gorovsky 1994; Mochizuki and Gorovsky 2004). MIC are transcriptionally silent in asexually dividing cells, featuring highly condensed chromatin and lacking transcription machinery (Fig. 1A). MIC differentiate into MAC during conjugation—the sexual phase of the *Tetrahymena* life cycle, accompanied by extensive genome rearrangements, which remove thousands of internally eliminated sequences (IES) and re-ligate the MAC-destined sequences (MDS) (Meyer and Chalker 2007). Transcription activation occurs in meiotic MIC and developing MAC, eventually establishing mRNA transcription in a euchromatin environment in MDS, while IES are destined for heterochromatin formation, and ultimately deletion, in a nuclear RNAi and *Polycomb* repression-dependent pathway (Mochizuki et al. 2002; Taverna et al. 2002; Liu et al. 2004; Liu et al. 2007).

**Figure 1.**
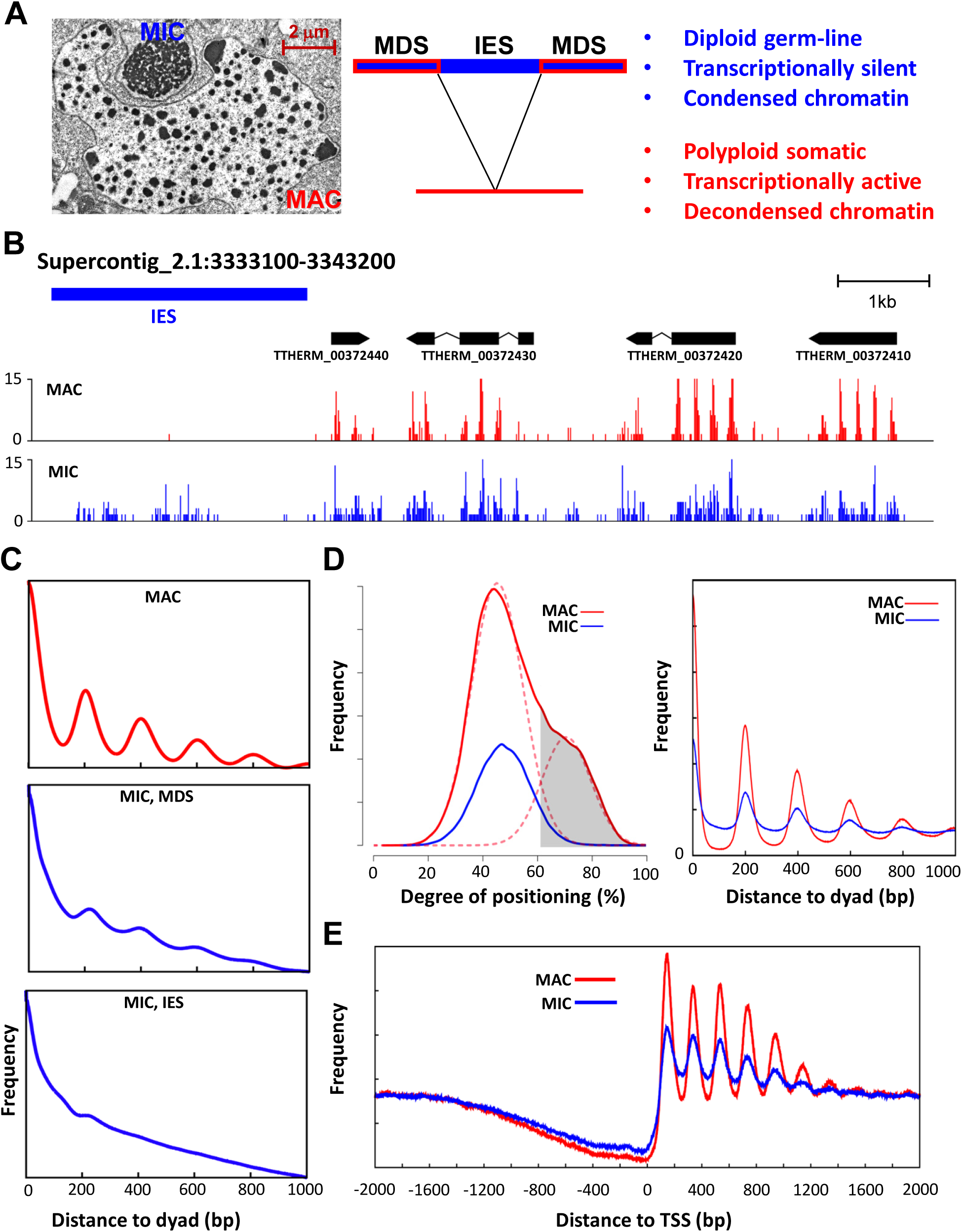
Nucleosome arrays are prominent in transcriptionally active MAC, but degenerate in transcriptionally silent MIC. A) Two types of structurally and functionally differentiated nuclei in *Tetrahymena*. Left, an electron micrograph showing the structurally differentiated macronucleus (MAC) and micronucleus (MIC), enclosed by their own nuclear envelope and contained in the same cytoplasmic compartment. Middle, DNA elimination accompanying MIC to MAC differentiation: IES, internal eliminated sequences, MIC-specific; MDS, MAC-destined sequences, shared between MAC and MIC. Right, functional differentiation of MAC and MIC. B) Nucleosome distribution in a representative genome region containing both MDS and IES, based upon MNase-Seq of the MAC and MIC samples. Paired-end MNase-Seq results from the MAC (red) and MIC (blue) samples are mapped to the MIC genome. Distribution of fragment centers representing nucleosome dyads is plotted, superimposed with models for gene bodies (black) and IES (blue). C) Phasogram of nucleosome distribution in MAC and MIC. x-axis: distance to fragment center (dyad); y-axis, frequency at a designated distance. MIC MNase-Seq results mapped to MDS and IES are plotted separately, while only MAC MNase-Seq results mapped to MDS (>99% of all mappable reads) is plotted. See Methods for details of calculations. D) Nucleosome delocalization in MIC compared with MAC. Left, nucleosome distribution according to their degrees of translational positioning, defined as the percentage of fragment centers within ±20 bp of the dyad of a called nucleosome, relative to the number of all fragment centers within the 147 bp nucleosome footprint. Only called nucleosomes supported by ≥50 fragments in the MAC sample are analyzed. The nucleosome distribution curve in MAC (red solid line) can be decomposed into two peaks of normal distribution (red dashed line), with the left peak representing more delocalized nucleosomes and the right peak well-positioned nucleosomes in MAC. Well-positioned nucleosomes in MAC (gray area, 61% cutoff) are selected for further analysis. They have significantly reduced degrees of translational positioning in MIC, as shown by a dramatic left shift. The new distribution (blue line) was similar to the more delocalized nucleosomes in MAC (red dashed line, left peak). See Methods for details and Supplemental File S1 for a compilation of properties of called nucleosomes. Right, composite distribution of fragment centers (dyad) from the MAC (red) and MIC (blue) samples, aligned to the dyads of the selected well-positioned nucleosomes (gray area in the left panel). E) Composite analysis of nucleosome positioning relative to TSS. Paired-end MNase-Seq results from the MAC (red) and MIC (blue) samples are mapped to the MAC genome. Distribution of fragment centers around TSS (± 2kb) is aggregated over 15,841 well-modeled genes. Note the stereotypical nucleosome arrays downstream of TSS. See Methods for details and Supplemental File S2 for a compilation of properties of well-modeled genes.

In this study, we employed MNase-Seq to reveal distinct nucleosome distribution patterns in structurally and functionally differentiated MAC and MIC of *Tetrahymena*. We found many well-positioned nucleosomes in MAC, downstream of TSS and flanking both the 5’ and 3’ splice sites in gene bodies. On the other hand, MIC nucleosomes were more delocalized, attributable to insufficiency in both nucleosome barriers and nucleosome dynamics conferred by transcription-associated trans-determinants. However, nucleosome occupancy in MIC and MAC were similar to each other, and fit predictions based upon DNA sequence features. In particular, arrays of well-positioned nucleosomes were often matched with oscillations in GC content, strongly supporting significant contributions from cis-determinants. We propose that cis- and trans-determinants may coordinately accommodate some well-positioned nucleosomes with important functions, driven by a positive feedback loop of the evolutionary processes, in which positioned nucleosomes shape the mutational landscape of associated DNA sequences, while the DNA sequences in turn reinforce nucleosome positioning. Our work provides a clear connection between nucleosome distribution and transcription, and lays the groundwork for future studies of nucleosome redistribution and *de novo* establishment of a transcription-compatible chromatin environment during MIC to MAC differentiation.

## Results

### Arrays of well-positioned nucleosomes are prominent in transcriptionally active MAC, but degenerate in transcriptionally silent MIC

We purified the structurally and functionally differentiated MAC and MIC from asexually dividing *Tetrahymena* cells (Fig. 1A). The mono-nucleosome was released by micrococcal nuclease (MNase) digestion, and the associated DNA was purified by agarose gel electrophoresis or sucrose gradient ultracentrifugation (Supplemental Fig. S1A-C). We performed paired-end Illumina sequencing of the mono-nucleosome DNA (MNase-Seq), and mapped the reads back to the *Tetrahymena* MAC (Eisen et al. 2006) and MIC reference genomes (*Tetrahymena* Comparative Sequencing Project, Broad Institute of Harvard and MIT: http://www.broadinstitute.org/annotation/genome/Tetrahymena/MultiHome.html). Here we focused mainly on the DNA samples prepared by heavy MNase digestion and agarose gel purification (referred to simply as the MAC and MIC samples, respectively; Supplemental Fig. S1D), due to their high recovery, low background, and unbiased representation. The *Tetrahymena* MIC genome contains ∽30% internally eliminated sequences (IES), with the rest being MAC-destined sequences (MDS). MNase-Seq reads from the MAC sample were virtually all mapped to the MAC genome or MDS in the MIC genome (>99%), confirming its high purity; while 26.6% of MIC MNase-Seq reads were mapped to MIC IES, close to the ∽30% estimate and consistent with only minor contamination of the MIC sample by MAC (Supplemental Fig. S1D).

In the MAC sample, well-positioned and regularly spaced nucleosomes were illustrated by the distribution of mapped MNase-Seq fragment centers—representing nucleosome dyads—in GBrowse views (Fig. 1B, MAC). Phasogram analysis of nucleosome positioning, measuring the distribution of fragment centers relative to one another, confirmed the presence of nucleosome arrays with ∽200 bp repeat length (NRL) and significant long-range order (Fig. 1C, MAC). We initially focused on translational nucleosome positioning, defined as the 147 bp genomic DNA occupied by a significant nucleosome sub-population and often represented by their exact dyad position. We calculated the degree of translational positioning for each called nucleosome in MAC (Fig. 1D, left panel: solid red line), defined as the number of fragment centers within ±20 bp of the called nucleosome dyad, relative to the number of all fragment centers within the 147 bp called nucleosome footprint (Zhang et al. 2009). The higher the number, the more definitive is the nucleosome positioning (with 100% corresponding to perfectly positioned nucleosomes, seldom reached due to imprecision in MNase-Seq mapping); the lower the number, the more delocalized is the nucleosome positioning (with ∽27% corresponding to random distribution). This analysis revealed a sub-population of well-positioned nucleosomes (Fig. 1D, left panel: dashed red line, right peak), featuring strong long-range order with ∽200 bp NRL (Fig. 1D, right panel). These nucleosomes were highly enriched within gene bodies (93.8%), which account for only ∽65% of the *Tetrahymena* MAC genome. Consistent with this observation, composite analysis of 15,841 well-modeled RNA polymerase II (Pol II) transcribed genes revealed prominent nucleosome arrays downstream of transcription start sites (TSS) (Fig. 1E).

The nucleosome distribution pattern observed in MAC, with strong translational positioning and long-range order, degenerated in MIC. This was illustrated by a side-by-side comparison in a representative genomic region of the nucleosome dyad distribution of the MAC and MIC samples (Fig. 1B). Phasogram analysis showed much reduced amplitudes of periodic nucleosome distribution in MDS (Fig. 1C, MIC MDS). The change was especially dramatic for IES, with only a small peak at ∽200 bp and no other detectable peaks further away from the composite nucleosome dyad (Fig. 1C, MIC IES). The result showed that nucleosome spacing was much more variable in MIC than MAC, even though the average NRL, as indicated by the peak positions, is still ∽200 bp (Fig. 1C). Compared with the well-positioned nucleosomes in MAC (Fig. 1D, left panel: red dashed line, right peak), degrees of translational positioning of their counterparts in MIC were dramatically reduced (Fig. 1D, left panel: blue line), to levels very similar to the delocalized nucleosomes in MAC (Fig. 1D, left panel: red dashed line, left peak). These nucleosomes also showed reduced long-range order in MIC relative to MAC (Fig. 1D, right panel), Composite analysis also revealed degeneration of nucleosome arrays in gene bodies in MIC (Fig. 1E). Our result strongly supports that nucleosomes in MIC are more delocalized relative to MAC, arguing that precisely phased nucleosome arrays in MAC are probably actively maintained by transcription-associated trans-determinants.

The phasogram analysis showed that attenuation of nucleosome periodicity was much more severe in IES than MDS (Fig. 1C). This interpretation was complicated by contaminating MAC in the MIC sample, aggravated by the high DNA content of polyploid MAC (∽45C) relative to diploid MIC (2C) as well as the high accessibility of the decondensed MAC chromatin relative to condensed MIC chromatin (Karrer 2012). Our results therefore only rigorously set the upper limit for the degree of translational positioning in MDS. Nonetheless, we argue that nucleosomes associated with IES—containing few, if any, normally expressed genes—are likely to be more delocalized than those associated with MDS. In other words, the IES result may set the lower limit for the degree of translational positioning in MDS.

We also studied nucleosome rotational positioning, using AAA/TTT distribution as a benchmark (Satchwell et al. 1986). Composite analysis of the trinucleotide distribution revealed periodic oscillations within the nucleosome footprint (Supplemental Fig. S2A). Spectrum analysis confirmed a 10.4 bp periodicity, with the peak intensity and position being essentially the same in both MAC and MIC (Supplemental Fig. S2B). This supports that rotational nucleosome positioning, in contrast to translational nucleosome positioning, is predominantly affected by DNA sequence features, consistent with observations in other eukaryotes (Albert et al. 2007; Mavrich et al. 2008; Brogaard et al. 2012; Cole et al. 2012).

### Nucleosome occupancy is strongly affected by cis-determinants

We calculated nucleosome occupancy—the probability for a particular DNA base pair to be associated with any nucleosomes—across *Tetrahymena* MAC and MIC genomes, based upon the MNase-Seq datasets. Despite substantial differences in nucleosome positioning in MAC and MIC (Fig. 1B), their nucleosome occupancy patterns were similar, as illustrated by a GBrowse view of the same genomic region (Fig. 2A). This was confirmed by a strong Spearman’s rank correlation (ρ=0.92) between the MAC and MIC MDS datasets. A probabilistic model, based upon the in vitro nucleosome distribution pattern in yeast (http://genie.weizmann.ac.il/software/nucleo_prediction.html) (Kaplan et al. 2009), fit well with nucleosome occupancy in both MAC (ρ=0.88) and MIC (ρ_IES_=0.85, ρ_MDS_=0.91). There was also a strong linear correlation between the predicted and measured nucleosome numbers in individual MAC scaffolds, MDS, and IES (Fig. 2B), supporting that at large distance scales (>1 kb), cis-determinants contribute uniformly to nucleosome occupancy in MIC and MAC. We further calculated the contribution from 30 individual DNA sequence features, including GC content (GC%), average slide and propeller twist, as well as occurrence of certain 3-mer and 4-mers (Fig. 2C and Supplemental Fig. S3) (Tillo and Hughes 2009; Wu et al. 2014). GC% showed strong positive correlations with the nucleosome occupancy in MAC (ρ=0.77) and MIC (ρ_IES_=0.84, ρ_MDS_=0.81), validating the widespread tracking behavior (Fig. 2A). Two structural features—average slide and propeller twist, scored for dinucleotide composition and correlated with GC%, were also strongly correlated with the nucleosome occupancy (Fig. 2C). On the other hand, WWW (W: A/T) showed strong negative correlation with the nucleosome occupancy in MAC (ρ=-0.70) and MIC (ρ_IES_=-0.75, ρ_MDS_=-0.84) (Fig. 2C). Significant negative correlations were also observed with the occurrences of various A/T-containing 4-mers (Supplemental Fig. S3). Both results were in line with reduced GC%, as well as the anti-nucleosomal nature of AT-rich tracts. Intriguingly, nucleosome occupancy in MIC, particularly IES, was even better correlated to many DNA feature-based predictions than MAC (Fig 2C and Supplemental Fig. S3). This suggests that nucleosome occupancy in MIC is more attributable to cis-determinants, while it is more perturbed in MAC by trans-determinants. We conclude that nucleosome occupancy is strongly affected by a limited number of cis-determinants, in particular GC% (Tillo and Hughes 2009).

**Figure 2.**
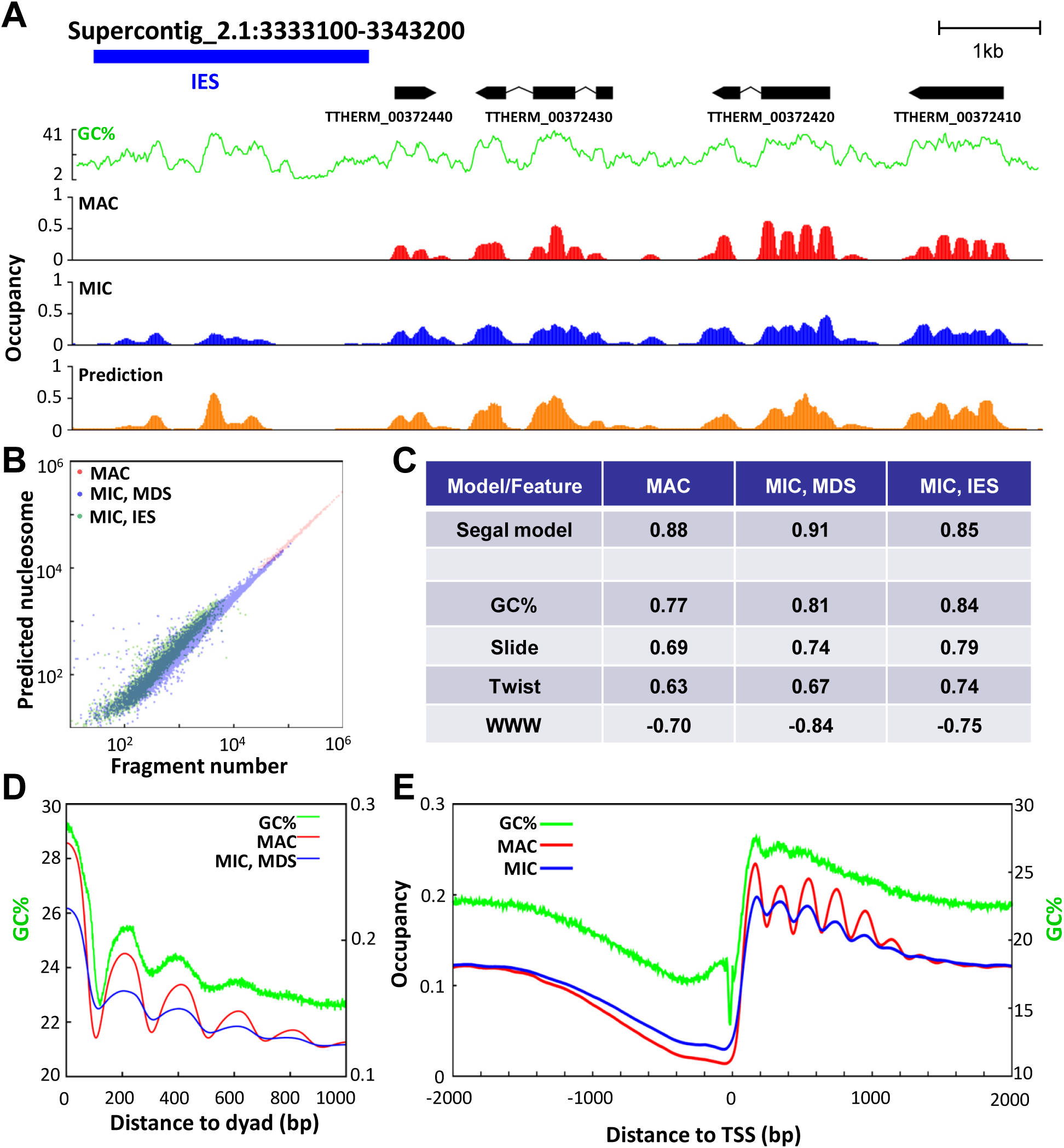
Nucleosome occupancy is strongly affected by cis-determinants. A) Nucleosome occupancy in a representative genomic region (as in Fig. 1B), based upon MNase-Seq of the MAC (red) and MIC samples (blue), as well as computational modeling (beige). GC% distribution (green) is superimposed. B) Strong correlations between nucleosome numbers measured by MNase-Seq and predicted by computational modeling over large distance scales. MAC: supercontigs in the MAC genome assembly; MIC MDS and IES: corresponding gap-free regions in the MIC genome assembly. See Methods for details and Supplemental File S3 and S4 for a compilation of IES and MDS, respectively. C) Spearman’s rank correlations between nucleosome occupancy levels, on the one hand calculated from the MAC and MIC MNase-Seq results, and on the other predicted by a comprehensive computational model (Segal model) or selected DNA sequence features (Kaplan et al. 2009; Tillo and Hughes 2009; Wu et al. 2014). The MIC MNase-Seq result is divided into MDS and IES. See Methods for details and Supplemental Fig. S3 for a complete list of 30 DNA sequence features. D) In phase oscillations of nucleosome occupancy and GC%. Nucleosome occupancy, based upon MNase-Seq of the MAC (red) and MIC samples (blue), is aligned to the dyads of the selected well-positioned nucleosomes (as in Fig. 1D). GC% distribution (green), also aligned to the dyads, is superimposed. E) Nucleosome occupancy relative to TSS. Nucleosome occupancy is aligned to and aggregated over the TSS of 15,841 well-modeled genes. GC% distribution (green), also aligned to TSS, is superimposed. Note the NFR upstream of TSS in the MAC (red) and MIC samples (blue), and the corresponding dip in GC%.

The relatively high GC% of nucleosomal DNA was consistent with, but not fully accounted for, by its enrichment in gene bodies (GC%: 22.3%), especially exonic regions (27.6%), and its depletion in inter-genic (17.8%) as well as intronic regions (16.3%). Composite analysis of nucleosome occupancy (aligned to the dyad of called nucleosomes) revealed the ∽200 bp nucleosome periodicity that degenerated in MIC (Fig. 2D), reminiscent of the phasogram analysis of nucleosome positioning (Fig. 1C). Composite analysis of GC% (also aligned to the dyad of called nucleosomes) showed that the core of nucleosomes featured significantly higher GC%, with sharp boundaries defined by precipitous transitions in GC% (Fig. 2D). Strikingly, there were significant GC% oscillations, in phase with ∽200 bp NRL in MAC and MIC (Fig. 2D). These results suggest that nucleosome occupancy is affected by periodic changes in GC%.

We further examined the nucleosome distribution in gene bodies. Composite analysis of all 15,841 well-modeled genes, aligned against TSS, showed only a small peak of GC% coincident with the +1 nucleosome position (Fig. 2E). We focused on well-positioned nucleosomes, which formed densely populated islands in nucleosome distribution maps plotted according to their distances from TSS (x-axis) and degrees of translational positioning (y-axis) (Fig. 3A: circled by dashed lines). Strikingly, composite analysis of genes containing the well-positioned +1 nucleosome revealed a prominent GC% peak, tracking the +1 nucleosome occupancy; composite analysis of genes containing the well-positioned +2 nucleosome revealed two prominent GC% peaks, tracking the +1 and +2 nucleosome occupancy; composite analysis of genes containing the well-positioned +3 nucleosome revealed three prominent GC% peaks, tracking the +1, +2, and +3 nucleosome occupancy (Fig. 3B). Furthermore, these well-positioned +1, +2, and +3 nucleosomes tended to be found together (Supplemental Fig. S4A), contained in high GC% peaks of gene bodies (Supplemental Fig. S4B). For genes with these well-positioned nucleosomes, the peak-to-trough differences in GC%, at more than 5%, were much higher than average, at less than 1% (Fig. 3B and Supplemental Fig. S4B). We conclude that GC% often oscillates in phase with occupancy of well-positioned nucleosomes, potentially playing a role in determining their distribution in gene bodies.

**Figure 3.**
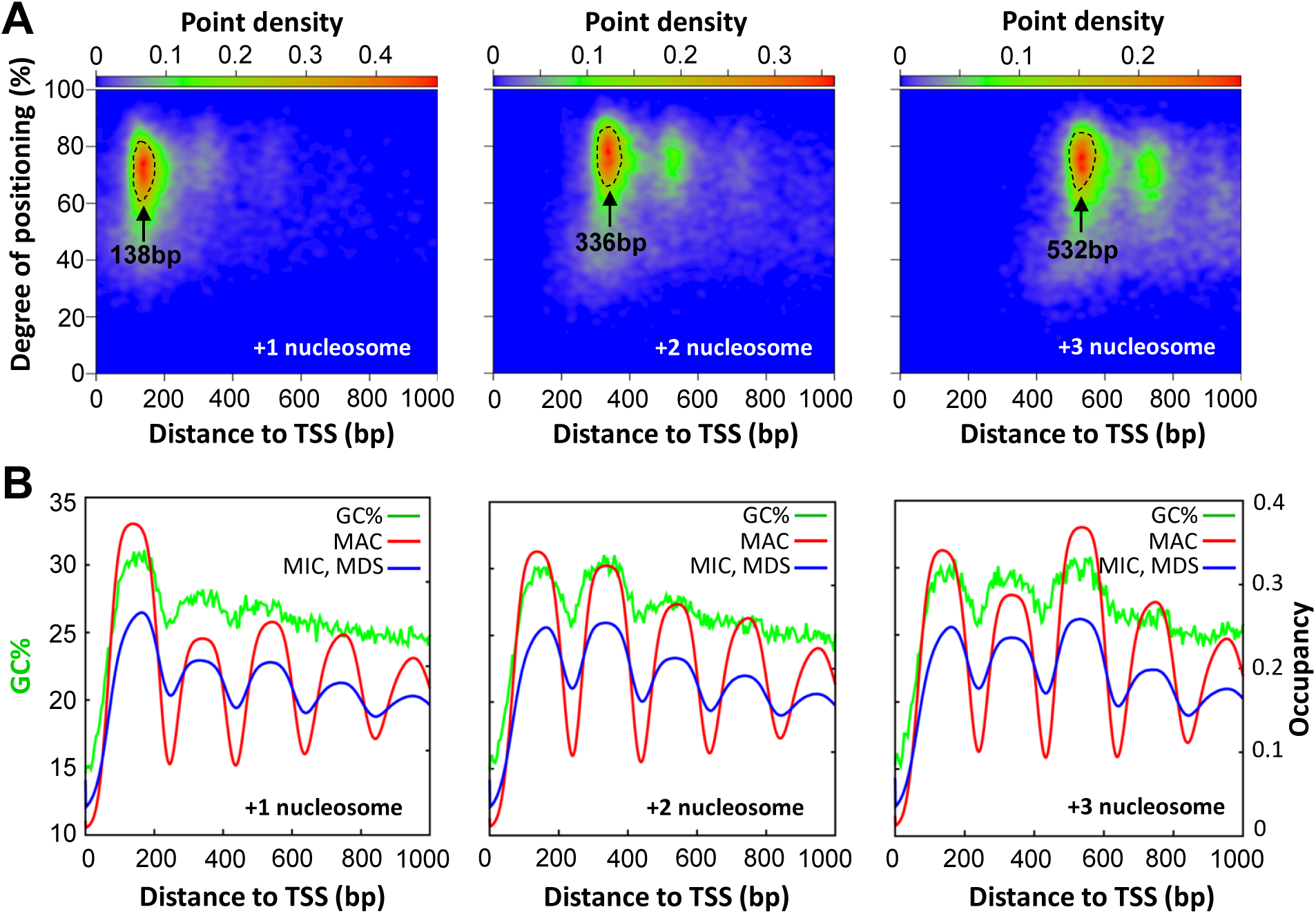
In phase oscillations of nucleosome occupancy and GC% downstream of TSS. A) Distribution of called nucleosomes in gene bodies (supported by ≥50 fragments in the MAC sample), according to their distances from TSS (x-axes) and degrees of translational positioning (y-axes). Note the clustering of nucleosome distributions, point density in color scales. Gene bodies containing these strongly clustered +1, +2, and +3 nucleosomes, their peak positions relative to TSS as indicated, are selected for further analysis (enclosed by dashed lines). See Methods for details and Supplemental File S1 for a compilation of properties of called nucleosomes. B) Composite analysis of nucleosome occupancy relative to TSS. MNase-Seq results from the MAC (red) and MIC (blue) samples are mapped to the MAC genome. Distribution of nucleosomes (occupancy) around TSS (-200 to +1000 bp) is aggregated over the genes containing the selected +1, +2, and +3 nucleosomes (enclosed by dashed lines in the left panels). GC% distribution (green), also aligned to TSS, is superimposed.

In both MAC and MIC, we also noted a very similar dip of nucleosome occupancy in nucleosome depleted regions (NDR) upstream of TSS and downstream of TES (Fig. 2E and Supplemental Fig. S5A). Composite analysis revealed significantly decreased GC%, almost parallel to nucleosome occupancy (Fig. 1E and Supplemental Fig. S5A). There was also accumulation of poly (dA:dT) tracts around TSS and TES (Fig. 4A and Supplemental Fig. S5B, C). MIC in asexually dividing cells contain no detectable levels of RNA polymerases (Mochizuki and Gorovsky 2004), transcription factors (Stargell and Gorovsky 1994), as well as transcription associated histone variants like H2A.Z (Stargell et al. 1993) and H3.3 (Cui et al. 2006). NDR in MIC, and to a large degree also in MAC, are therefore independent of these trans-determinants, and are instead underlain by low GC% and prevalence of anti-nucleosomal sequences, consistent with observations of in vitro nucleosome reconstitution using yeast genomic DNA (Kaplan et al. 2009; Zhang et al. 2009). Taken together with the in phase distribution of nucleosome arrays and GC% in gene bodies, our results support that nucleosome occupancy is strongly affected by cis-determinants.

**Figure 4.**
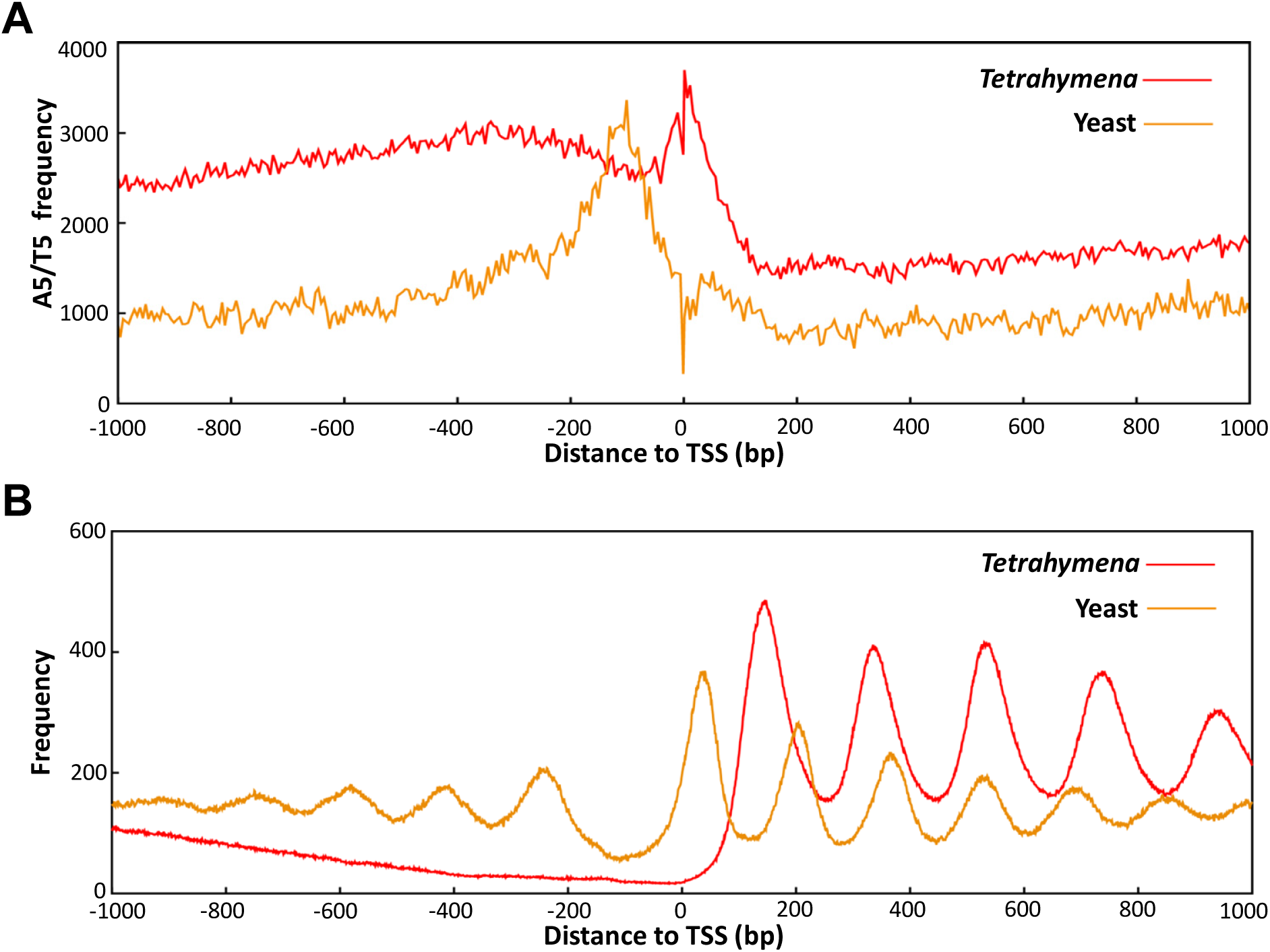
The +1 nucleosome placement is further downstream of TSS in *Tetrahymena* than in yeast. A) Poly (dA:dT) tract distribution relative to TSS in *Tetrahymena* (red) and yeast (beige). A5/T5 (AAAAA or TTTTT) and A7/T7 distributions are calculated around TSS (± 1kb) using the 5 bp bin. B) Nucleosome distribution relative to TSS in *Tetrahymena* (red) and yeast (beige). Paired-end MNase-Seq results from *Tetrahymena* MAC (Fig. 1 and Supplemental Fig. S1) and yeast (Kent et al. 2011) are analyzed. Distribution of fragment centers around TSS (± 1kb) is aggregated over 15,841 genes in *Tetrahymena* (Supplemental File S2) and 2,231 genes in yeast (Zhang and Dietrich 2005). Note the downstream placement of the +1 nucleosome (Yeast: 56 bp; *Tetrahymena*: 138 bp), increase in NRL (Yeast: 167 bp; *Tetrahymena*: 200 bp), and lack of nucleosome arrays upstream of TSS, when *Tetrahymena* is compared with yeast. See Methods for details.

### The +1 nucleosome placement is further downstream of TSS in Tetrahymena than in yeast

Unlike yeast (Albert et al. 2007), the +1 nucleosome in *Tetrahymena* was separated from TSS by a large distance, as best illustrated in the nucleosome distribution plotted according to their distances from TSS and degrees of translational positioning (Fig. 3A). The heat maps highlighted a strong tendency for the +1 nucleosomes with high degrees of translational positioning to cluster in a narrow band downstream of TSS with nucleosome density peaking at 138 bp (Fig. 3A). The +2 and +3 nucleosomes with high degrees of translational positioning clustered at around 336 bp and 532 bp (Fig. 3A), consistent with the +1 nucleosome placement ∽138 bp downstream of TSS and NRL of ∽200 bp (nucleosome distances to TSS: 138+200n bp, n=0, 1, 2 …). Nucleotide composition anomalies were observed in TSS (Supplemental Fig. S6A) as well as TES (Supplemental Fig. S6B). These anomalies may directly affect nucleosome distribution through cis-determinants. In apparent contrast to yeast, poly (dA:dT) tracts were highly enriched around TSS in *Tetrahymena* (Fig. 4A), incompatible with nearby placement of the +1 nucleosome. Alternatively but not mutually exclusively, these anomalies may represent DNA sequence motifs for recruiting other chromatin-associated proteins, thus indirectly affecting nucleosome distribution through trans-determinants. In support, the TATA box was enriched around TSS (Supplemental Fig. S6C), while the termination signal (AATAAA) was enriched near TES (Supplemental Fig. S6D). We conclude that the +1 nucleosome placement is further downstream of TSS in *Tetrahymena* than in yeast (Fig. 4B). The downstream placement of the +1 nucleosome is more commonly observed in metazoa, associated with promoter proximal pausing of Pol II (Kwak and Lis 2013). This implies that as a mechanism for transcription regulation, promoter proximal pausing may have arisen early in eukaryotic evolution.

### Nucleosome arrays in gene bodies are affected by transcription levels

The stereotypical nucleosome arrays generally became less prominent with decreasing expression levels of associated genes (Fig. 5A and Supplemental Fig. S7A, B: divided into 10 quantiles by their expression ranking in asexually dividing cells), reflected most prominently by the changing amplitudes of the +1, +2, and +3 nucleosomes. Indeed, there was dramatic and progressive degeneration of the nucleosome arrays in genes with little or no expression, corresponding to the last three quantiles (Fig. 5A and Supplemental Fig. S7A, B). This was mainly attributed to reduced degrees of translational positioning (Fig. 5B, Supplemental Fig. S7C). Intriguingly, the result suggests a shift from the well-positioned nucleosomes to the delocalized nucleosomes, as transcription activity ceases. The result was reminiscent of the shift from the well-positioned nucleosomes in MAC to the delocalized nucleosomes in MIC, in the presence and absence of transcription, respectively (Supplemental Fig. S7D). Indeed, the numbers of well-positioned +1, +2, and +3 nucleosomes decreased significantly and monotonically in the last three quantiles of genes (Supplemental Fig. S7E). To a lesser degree, nucleosome arrays were also disrupted in genes with the highest expression levels in vegetatively growing cells (Supplemental Fig. S8A-D). In an extreme case, there were no positioned nucleosomes in the highly transcribed region of the rDNA mini-chromosome in MAC (Supplemental Fig. S8E). Taken together, our results strongly support that the transcription process significantly affects the stereotypical nucleosome arrays, which degenerate at very low rates of transcription, and are disrupted at very high rates.

**Figure 5.**
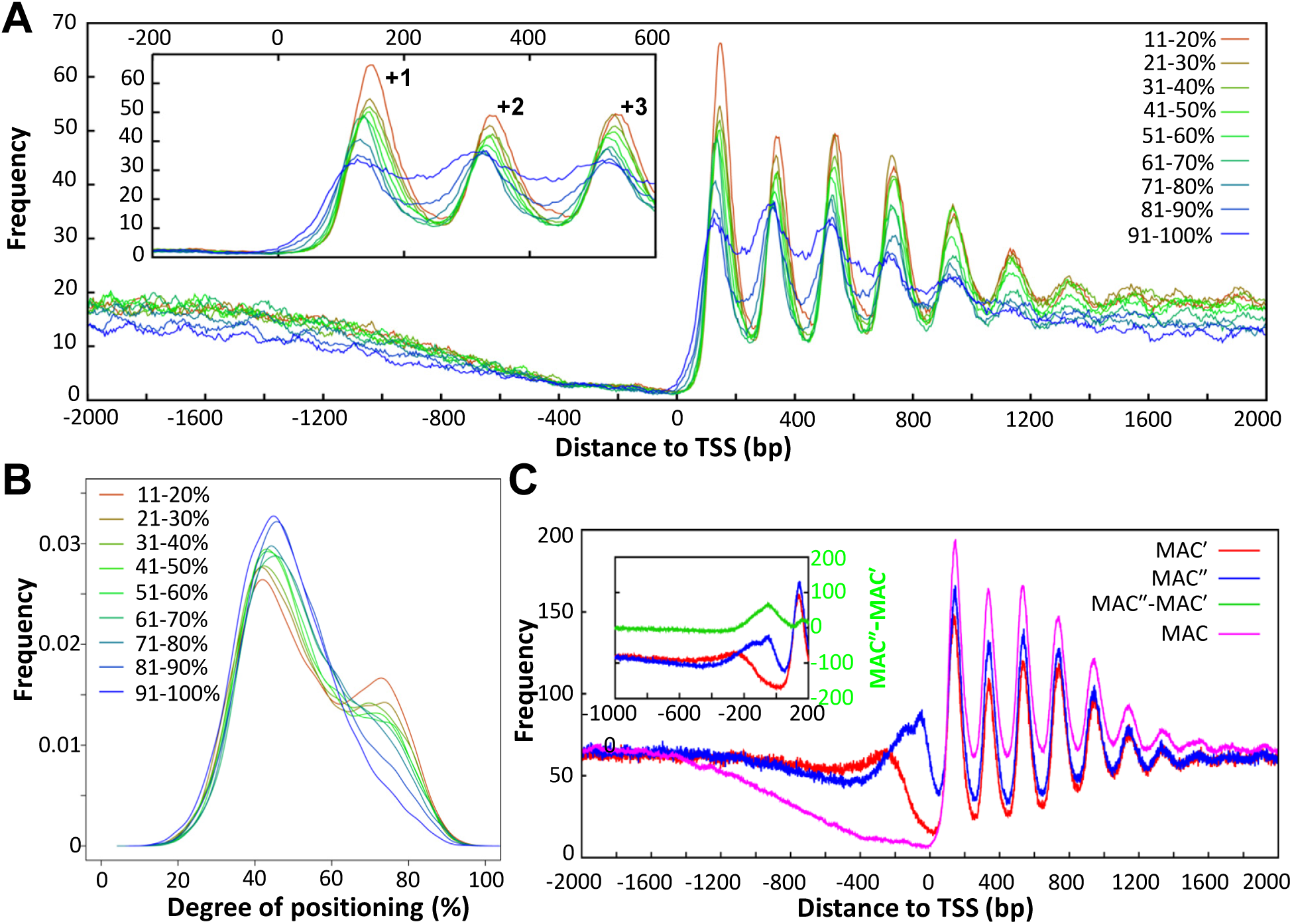
Transcription-associated trans-determinants affect nucleosome positioning in gene bodies. A) Stereotypical nucleosome arrays in genes with different transcription levels. 15,841 well-modeled genes are ranked from high to low by their expression levels and divided into 10 quantiles. Distribution of MAC MNase-Seq fragment centers around TSS is aggregated over all genes in a quantile. Distributions for quantile 2-10, normalized to per million reads per gene, are plotted. Inset: zoom in on the +1, +2, and +3 nucleosomes. Note that the amplitudes of the periodic nucleosome distribution decrease monotonically with decreasing expression levels. See Supplemental Fig. S7 for more details. B) Translational nucleosome positioning in genes with different transcription levels. Nucleosome distribution is aggregated over all called nucleosomes in gene bodies of a specified quantile. Normalized distributions for quantile 2-10 are plotted. Note that as transcription levels decrease, degrees of translational positioning of associated nucleosomes also decrease, with reduction in the well-positioned nucleosome peak and increase in the delocalized nucleosome peak. C) Comparison of the nucleosome distribution patterns around TSS from three differently prepared MAC samples. Inset: the difference between the MAC′ and MAC″ samples, highlighting the non-nucleosomal footprint near TSS (green).

**Figure 6.**
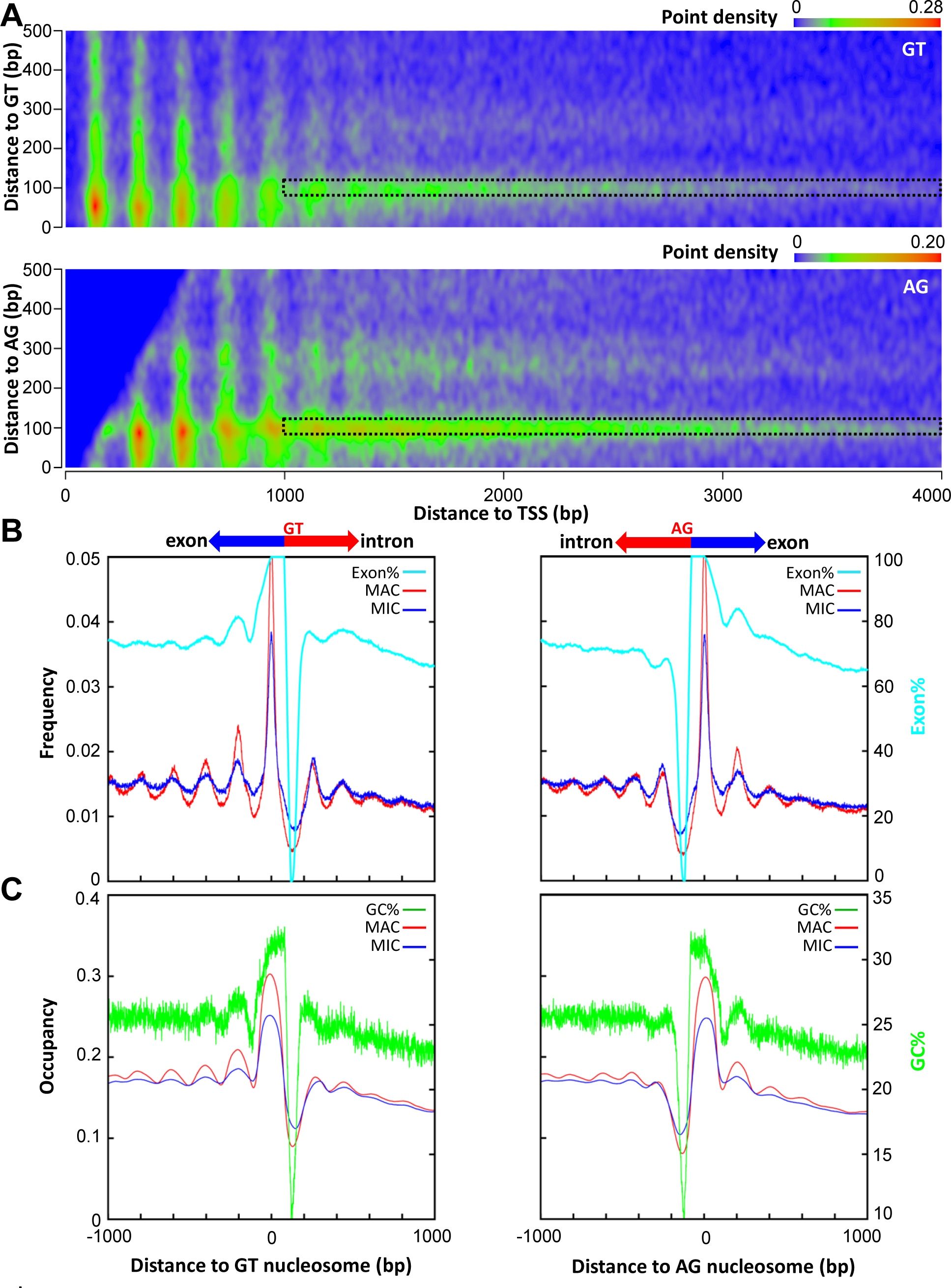
Splice sites are flanked by positioned nucleosomes. A) Distribution of called nucleosomes in gene bodies (supported by ≥50 fragments in the MAC sample), according to their distances from TSS (x-axes) and splice sites (y-axes; top: GT; bottom: AG). Note the clustering of nucleosome distribution along both x- and y-axis, point density in color scales. Nucleosomes predominantly defined by their positions relative to splice sites (1000≤x≤4000, 80≤y≤120) are selected for further analysis (enclosed by dashed lines). See Supplemental File S1 for a compilation of properties of called nucleosomes. B) Composite distribution of fragment centers from the MAC (red) and MIC (blue) samples, aligned to the dyads of the called nucleosomes flanking splice sites (left: GT, right: AG). The frequency value is normalized to per million reads per nucleosome. The distribution of exon% (pale blue), calculated as the probability that a particular position containing exons, is superimposed. Note that the propagation of nucleosome arrays near splice sites is accompanied by the in phase oscillation of exon%. See Methods for details. C) Composite distribution of nucleosome occupancy from the MAC (red) and MIC (blue) samples, aligned to the dyads of the called nucleosomes flanking splice sites (left: GT, right: AG). The distribution of GC% (green) is superimposed. Note the correlation between nucleosome arrays near splice sites and the oscillation of GC%.

We also compared the nucleosome distribution patterns around TSS from three differently prepared MAC samples (Fig. 5C). In addition to the sample generated by heavy MNase digestion, we also included two samples generated by light MNase digestion (Supplemental Fig. S1), which recovered more MNase-sensitive chromatin at inter-genic regions (Fig. 5C). One sample was derived from the mono-nucleosome fractions of the soluble chromatin after sucrose gradient ultracentrifugation (referred to as MAC′), the other was derived from gel purified mono-nucleosome sized DNA fragments (MAC″) (Supplemental Fig. S1). MAC′ was very selective for the mono-nucleosome; while MAC″ contained DNA associated with other chromatin bound proteins. Their MNase-Seq patterns around and particularly upstream of TSS were significantly different from each other and from that of heavy MNase digestion: MAC′ revealed the -1 nucleosome, distributed in a broad band 200-400 bp upstream of TSS; while MAC″ revealed a distinct set of peaks much closer to TSS, dominated by a non-nucleosomal footprint (Fig. 4C). Our interpretation was further corroborated by comparing the MNase-Seq data from gene quantiles of different expression levels. With decreasing levels of gene expression, the NDR between the -1 and +1 nucleosome showed increasing levels of nucleosome occupancy in MAC′ (Supplemental Fig. S7A), arguing for promoter occlusion by labile nucleosomes (Jin et al. 2009; Weiner et al. 2010; Xi et al. 2011); while the non-nucleosomal footprint dwindled to background levels in MAC″ (Supplemental Fig. S7B), indicative of reduced promoter binding. The MNase-Seq patterns upstream of TSS were therefore correlated with the transcription levels in opposite ways in MAC′ and MAC″, supporting competitive binding of labile nucleosomes and non-nucleosomal proteins (Workman and Kingston 1992). The latter is attributable to transcription-associated trans-determinants, which may include specific transcription factors, the general transcription machinery, and ATP-dependent chromatin remodelers (Henikoff et al. 2011; Kent et al. 2011). Together with the cis-determinants, these trans-determinants may act like nucleosome barriers to shape the chromatin environment around TSS, or provide energy input for maintaining the stereotypical nucleosome arrays in gene bodies.

### Splice sites are flanked by positioned nucleosomes

*Tetrahymena* contain an average of 3.61 introns per gene, with introns significantly shorter than exons on average (intron: 141 bp; exon: 420 bp). Internal exons had much higher levels of nucleosome occupancy than the flanking intronic regions in both MAC and MIC (Supplemental Fig. S9A), consistent with GC% being much higher in exons (27.6%) than introns (16.3%) (Supplemental Fig. S9B). This distribution is similar to that in metazoa and plants (Schwartz et al. 2009; Tilgner et al. 2009; Chodavarapu et al. 2010). Composite analysis revealed positioned nucleosomes in exonic regions flanking the 5’ (GT) splice site and, even more prominently, 3’ (AG) splice sites (Supplemental Fig. S9C, D). To further dissect the contributions of TSS and splice sites, we plotted heat maps of nucleosomes distributed at different distances from these two landmarks (Fig. 5A). As expected, nucleosomes positioned relative to TSS manifested as a set of evenly spaced vertical streaks, which dominated the left side of the plots but diminished as the distances from TSS increased (Fig. 5A). A conspicuous horizontal streak, representing nucleosomes positioned at ∽100 bp from the splice sites, was also revealed (Fig. 5A). It persisted much further into gene bodies, and dominated the right side of the plots (Fig. 5A). This grid-like pattern still held, even when we focused on nucleosomes with high levels of occupancy and degrees of translational positioning (Supplemental Fig. S9E). Composite analysis of nucleosomes dominantly affected by splice sites revealed that they were strongly positioned in MAC, and their positioning was only moderately reduced in MIC (Fig. 5B). Composite analysis also revealed that the occupancy levels of these nucleosomes coincided with a prominent GC% peak in exons, right next to a stretch of DNA with very low GC% at the intron side (Fig. 5C). The presence of these strong anti-nucleosomal sequences may serve as a barrier, against which nucleosomes in exons are positioned. However, the positioning of neighboring nucleosomes degenerated quickly away from splice sites (Fig. 5B), in apparent contrast with the long range order in stereotypical nucleosome arrays (Fig. 1E), most likely maintained by ATP-dependent chromatin remodelers (Zhang et al. 2011; Yen et al. 2012). Furthermore, compared with the stereotypical nucleosome arrays (Fig. 4A and Supplemental Fig. S7A, B), nucleosomes positioned next to splice sites were not much affected by transcription levels (Supplemental Fig. S9D). All this suggests that transcription-associated trans-determinants do not affect nucleosome distribution around splice sites as much as they do downstream of TSS. We conclude that splice sites are flanked by positioned nucleosomes, and contribute to the chromatin environment of gene bodies independent of TSS. These positioned nucleosomes may potentially provide additional information to allow precise recognition of splice sites, or even modulate alternative splicing.

## Discussion

We have demonstrated distinct nucleosome distribution patterns in MIC and MAC, two structurally and functionally differentiated nuclei in a unicellular eukaryote, *Tetrahymena thermophila*. Arrays of well-positioned nucleosomes coincide with active transcription in MAC, while their degeneracy is associated with transcriptional silencing in MIC. This is mainly attributed to nucleosome delocalization in MIC, while in MAC, nucleosome arrays are maintained by barriers comprised of trans-as well as cis-determinants, and energy input from transcription-coupled events. All this is consistent with positive feedback to maintain the proper chromatin environment for transcription, potentially involving specific transcription factors, the general transcription machinery, as well as accompanying ATP-dependent chromatin remodelers. As a manifestation of nuclear dimorphism, MAC features a chromatin environment predominantly shaped by active transcription, likely regulated by promoter proximal pausing of Pol II, while nucleosome distribution in MIC is more dictated by cis-determinants in the absence of transcription. Importantly, our work also sets the stage for future studies of nucleosome redistribution and de novo establishment of the transcription-compatible chromatin environment during MIC to MAC differentiation, allowing us to address key epigenetic questions in an experimentally tractable system.

### Arrays of well-positioned nucleosomes in MAC but not MIC

We compared the nucleosome distribution patterns in MAC and MIC. In the transcriptionally active MAC, arrays of well-positioned nucleosomes were prominent in gene bodies, especially downstream of TSS. They were consistently revealed by different procedures for preparing MNase-Seq samples. As this stereotypical nucleosome distribution pattern has also been found in yeast and other eukaryotes, it strongly suggests that nucleosome organization is dependent upon general, if not universal, mechanisms (Struhl and Segal 2013). In the transcriptionally silent MIC, these nucleosome arrays degenerated, as shown by the dramatic reduction in translational nucleosome positioning and long-range order. This strongly suggests that the precise distribution pattern in MAC is actively maintained by transcription-associated trans-determinants missing in MIC. Conversely, the transcription process in MAC may be regulated by its nucleosome distribution pattern, as the downstream placement of the +1 nucleosome is implicated in promoter proximal pausing of Pol II. Well-positioned nucleosomes were also found in MAC in contexts other than transcription, in regions adjoining splice sites (Fig. 6), replication origins, and the telomere of the rDNA mini-chromosome (Supplemental Fig. S10). Our data and interpretation are consistent with the barrier-diffusion model, in which an anti-nucleosomal sequence and/or strong binding of sequence-specific factors generate a nucleosome barrier and subsequently establish neighboring phased arrays by statistical positioning and/or chromatin remodeling (Kornberg and Stryer 1988; Mobius et al. 2013).

Nucleosome delocalization in MIC may be caused by highly variable nucleosome spacing, due to reduced ATP-dependent chromatin remodeler activities, which are often associated with transcription and play a significant role in shaping nucleosome arrays in gene bodies (Yen et al. 2012). Alternatively, it may be attributed to shifting of nucleosome arrays in either the upstream or downstream direction in individual cells, due to insufficiency of nucleosome barriers, which are often defined by binding proteins including the general transcription machinery and specific transcription factors (Rhee and Pugh 2011; Rhee and Pugh 2012). Lack of strong phasing in our MIC dataset is consistent with both possibilities, and underlies the weak signal corresponding to ∽200 bp NRL. More accurate and precise measurement of nucleosome spacing in MIC will require highly purified MIC, as well as increased mapping resolution, possibly by adopting hydroxyl radical cleavage (Brogaard et al. 2012).

### Placement of the +1 nucleosome is connected with promoter proximal pausing of Pol II

The stereotypical nucleosome arrays downstream of TSS shape the epigenetic landscape for Pol II transcription (Hughes and Rando 2014). The +1 nucleosome in *Tetrahymena* was generally placed much further away from TSS than in yeast, in positions similar to metazoa. In yeast, gene promoters are often wrapped up in the +1 nucleosome, directly implicating it in the regulation of transcription initiation (Lieleg et al. 2014). In metazoa, transcription initiation and promoter proximal pausing are regulated separately, with the latter affected by the +1 nucleosome (Kwak et al. 2013; Kwak and Lis 2013). Pol II pausing may also directly impact on the +1 nucleosome positioning, forming a feedback loop to maintain the optimal chromatin environment for transcription regulation. In certain unicellular eukaryotes (e.g., *Dictyostelium*) as well as metazoa (Chang et al. 2012), promoter proximal pausing is generally assumed to be caused by the negative elongation factor (NELF), a multi-protein complex shared among metazoa (Kwak and Lis 2013). However, similar patterns have also been observed in *Arabidopsis* (Li et al. 2014) as well as *Tetrahymena*, which have no apparent NELF homologues. This strongly suggests that there is an evolutionarily ancient, NELF-independent mechanism for promoter proximal pausing of Pol II, although it remains a formal possibility that protozoa and plants may feature NELF homologues too remotely related with their metazoan counterparts to be identified by direct sequence comparison.

### Cis- and trans-determinants coordinately accommodate well-positioned nucleosomes

The strong correlation between the nucleosome occupancy levels in MAC and MIC, featuring distinct trans-determinants, demonstrated a prominent role played by cis-determinants. This was corroborated by computational modeling, which predicted the nucleosome occupancy based upon *Tetrahymena* DNA sequence features. Detailed analysis further revealed that GC content was the dominant cis-determinant for nucleosome distribution. The apparent discrepancy between the distinct nucleosome positioning and highly correlated nucleosome occupancy in MAC and MIC can be reconciled if nucleosome distribution in MIC is a more delocalized form of the MAC distribution. Specifically, in both MAC and MIC, nucleosomes are enriched in high GC% sequences and distributed in degenerate sites with ∽10 bp intervals to conserve rotational positioning. However, nucleosomes may be more narrowly distributed around their average positions in MAC, while more spread out in degenerate sites in MIC. Given the length of *Tetrahymena* linker DNA (>50 bp), it is conceivable that active participation of trans-determinants, particularly energy input from ATP-dependent chromatin remodelers, may be needed to maintain uniform NRL (Budarf and Blackburn 1986; Tsankov et al. 2010; Zhang et al. 2011; Hughes et al. 2012). Indeed, computational modeling based upon DNA sequence features consistently predicted the nucleosome occupancy in MIC MDS with better scores than in MAC, arguing for significant perturbations from trans-determinants in the latter.

Nucleosome positioning may be anchored by juxtaposition of anti-nucleosomal and energetically favorable sequences, exemplified by nucleosomes associated with container sites in human cells (Valouev et al. 2011). What is striking in *Tetrahymena* MAC is that oscillations in GC% and nucleosome occupancy often remain in phase beyond the distance covered by a single nucleosome, especially in the stereotypical nucleosome arrays downstream of TSS. Combined with the 10.4 bp periodicity underlying nucleosome rotational positioning, our results suggest that *Tetrahymena* genome sequence may accommodate, in coordination with trans-determinants, well-positioned nucleosomes in MAC. Importantly, well-positioned nucleosomes may confer advantages in natural selection, as transcription initiation from cryptic sites occurs when regular nucleosome arrays in gene bodies are perturbed (Whitehouse et al. 2007).

It has recently emerged that positioned nucleosomes may affect the mutational landscape across the genome (Sasaki et al. 2009; Prendergast and Semple 2011). Nucleosomes function as barriers to DNA replication in the lagging strand, with their centers coinciding with the 5’ end of Okazaki fragments (Smith and Whitehouse 2012), which are synthesized by the error-prone DNA polymerase α and retained in a substantial amount in the replicated genome (Reijns et al. 2015). This leads to significantly elevated nucleotide substitution rates around nucleosome dyads (Sasaki et al. 2009; Prendergast and Semple 2011), providing mutations to drive evolution. Nucleosome occupancy may even alter mutation spectra, leading directly to higher GC% (Prendergast and Semple 2011; Chen et al. 2012). We propose that cis- and trans-determinants may coordinately accommodate some nucleosomes with important functions, driven by a positive feedback loop of the evolutionary processes, in which positioned nucleosomes shape the mutational landscape of associated DNA sequences, while the DNA sequences in turn reinforce nucleosome positioning (Fig. 7).

**Figure 7.**
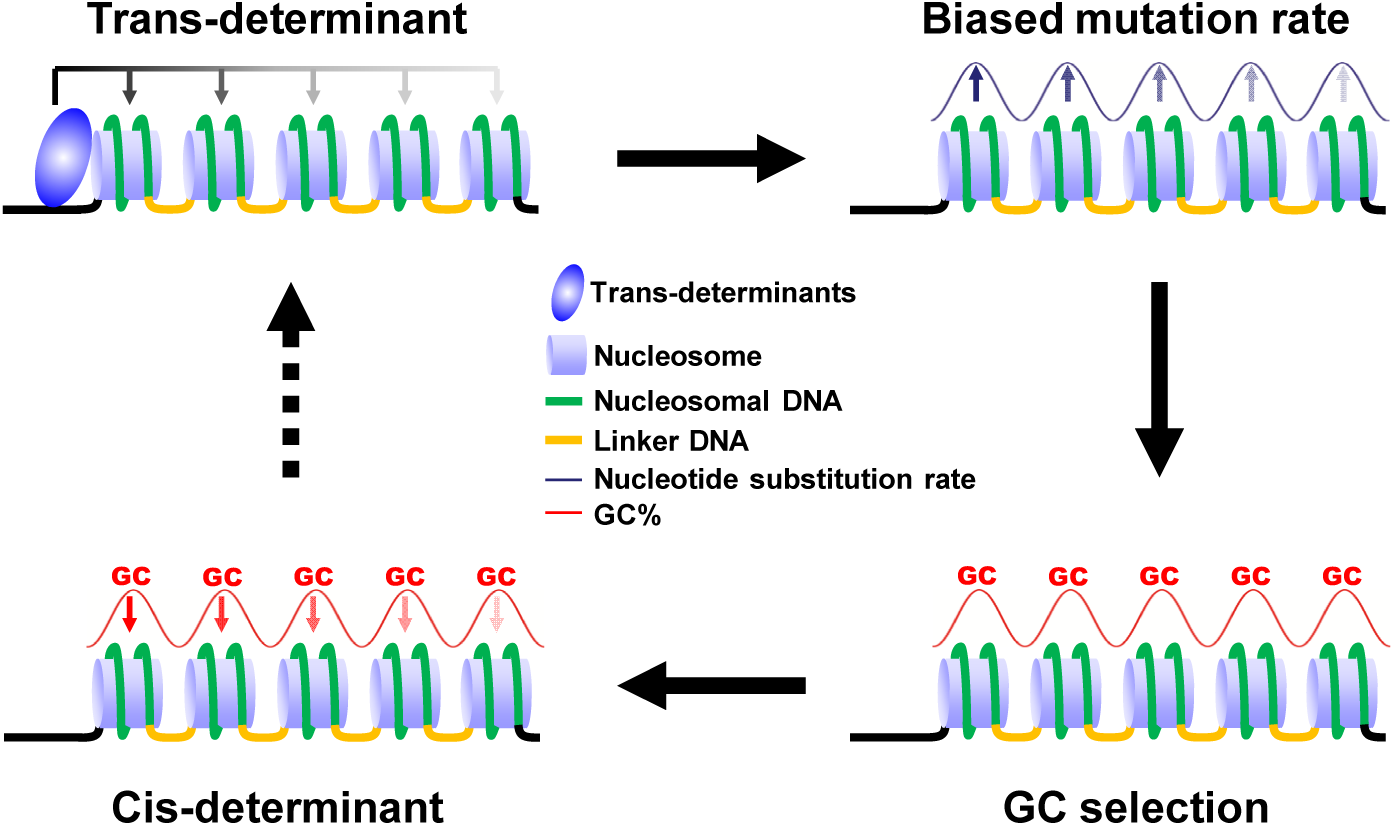
A positive feedback allows cis- and trans-determinants to coordinately accommodate well-positioned nucleosomes. See text for details.

## Methods

Additional details are available in the Supplemental Methods.

### Cell culture, purification of nuclei, and MNase digestion

*Tetrahymena thermophila* wild-type strain CU428, obtained from the *Tetrahymena* Stock Center (https://tetrahymena.vet.cornell.edu/), was grown in SPP medium at 30 °C (Gorovsky et al. 1975). After cell lysis, nuclei were isolated by differential centrifugation following established protocols (Sweet and Allis 2006; Papazyan et al. 2014). MAC fractions containing less than 5% MIC (count/count) and MIC fractions containing less than 0.1% MAC (count/count) were used for MNase digestion. MNase digestion was carried out at 25 °C for 15 min (50 mM Tris-HCl (pH 8), 5 mM CaCl_2_, 1mM β-mercaptoethanol, 0.1% NP-40, 0.1mg/ml BSA, 20 U (for light digestion) or 100 U MNase (for heavy digestion)), stopped by adding 10 mM EGTA and 1 mM EDTA. Mono-nucleosome sized DNA was selected by agarose gel purification; alternatively, mono-nucleosomes were purified by sucrose gradient ultracentrifugation.

### MNase-Seq data processing

We performed paired-end Illumina sequencing of mono-nucleosomal DNA (MNase-Seq), and mapped the reads back to the *Tetrahymena* reference genome assemblies using TopHat2 (Trapnell et al. 2012). The MAC reference is from the *Tetrahymena* genome database (TGD: http://ciliate.org) (Eisen et al. 2006; Stover et al. 2012); the MIC reference is from the *Tetrahymena* Comparative Sequencing Project (http://www.broadinstitute.org/annotation/genome/Tetrahymena/MultiHome.html). Unique mapped reads were extracted, and only one of any potential PCR duplicates (fragments, defined by a pair of properly mapped reads, with the same location in the genome) were kept. Only fragments ranging from 120 to 260 bp in length were analyzed. The mapping results were visualized using GBrowse 2.0 (Stein 2013).

### Nucleosome calling and analyses of translational nucleosome positioning

Nucleosomes were called using the NucPosSimulator (Schopflin et al. 2013), which identify non-overlapping nucleosomes using MNase-Seq fragment centers. We calculated the degree of translational positioning for each called nucleosome in MAC, defined as the number of fragment centers within ±20 bp of the called nucleosome dyad, relative to the number of all fragment centers within the 147 bp called nucleosome footprint (Zhang et al. 2009). The MAC nucleosome distribution curve, plotted according to their degrees of translational positioning, was decomposed into two Gaussian distribution components, using the mixtools package in R (Benaglia et al. 2009).

### Phasogram and spectrum analysis

Periodicity in the MNase-Seq data was identified by phasogram (Valouev et al. 2011), aligned to and aggregated for all fragment centers across the genome. Only one side was plotted, due to symmetry. For rotational positioning, we focused on the distribution of the AAA/TTT trinucleotide instead of the AA/TT dinucleotide, to reduce the background noise due to the low GC% of the *Tetrahymena* genome. Composite frequency of the AAA/TTT trinucleotide in the nucleosomal DNA was calculated and aggregated in the 1 kb region downstream of all aligned fragments ends. At each position i in the alignment, the percentage of AAA/TTT trinucleotide at the [i-1, i, i+1] positions was calculated. The periodicity of the AAA/TTT trinucleotide was identified by spectrum analysis with discrete Fourier transform, using the “ spectrum” function in R (smoothing width: 9) (http://www.R-project.org).

### Nucleosome distribution relative to TSS, TES, and splice sites

For analyses related to TSS, TES, and splice sites, we focused on 15,841 gene models strongly supported by RNA-Seq results (Xiong et al. 2012), hereafter referred to as well-modeled genes (see Supplemental Methods for details). Selected TSS and TES are supported by at least 5 independent RNA-Seq reads. Selected splice sites are supported by at least 1 RNA-Seq read spanning the intron-exon junction. Composite analyses of MNase-Seq fragment centers, nucleosome occupancy, and called nucleosome dyads were performed after aligning to the selected landmarks.

### Ranking of genes according to their expression levels

Expression levels of 15,841 well-modeled genes in asexually dividing cells—from which the MNase-Seq samples were prepared—were quantified using the RNA-Seq data (Xiong et al. 2012). Reads were mapped using Tophat2 (Trapnell et al. 2012), and the fragments per kilobase of exon per million mapped fragments (FPKM) values were generated using the Cuffdiff program (Trapnell et al. 2012). Genes were ranked from high to low by their FPKM values and partitioned to quantiles accordingly.

### Nucleosome occupancy from the MNase-Seq datasets or DNA sequence features

A uniform 147 bp nucleosomal footprint was assumed for all MNase-Seq fragments, extending 73 bp in both directions from their centers. Coverage for each genomic position was calculated based on these 147 bp footprints. Computational predictions were made using a comprehensive model built upon a dataset from nucleosomes reconstituted with yeast histones and genomic DNA in vitro (Kaplan et al. 2009). Software for this model was downloaded from the Segal Lab website (http://genie.weizmann.ac.il/software/nucleo_prediction.html). Computational predictions were also made using the 14 DNA sequence features selected for their dominant effects on nucleosome occupancy (Tillo and Hughes 2009). The other 16 features were created for a model based on geometrically transformed Tsallis entropy (Wu et al. 2014). Spearman rank correlations were calculated to compare nucleosome occupancy in matching sequences in the MAC and MIC, computational predictions to nucleosome occupancy, and feature values to nucleosome occupancy.

### Nucleosome distribution heatmap

Nucleosome distribution heatmaps were plotted according to: 1) distances of called nucleosome dyads to TSS (x-axis) and degrees of translational positioning (y-axis); 2) distances of called nucleosome dyads to TSS (x-axis) and splice sites (y-axis), using the Spatstat package in R (smoothing parameter sigma: 10) (Baddeley and Turner 2005). Point densities— referred to as z values below—were represented in color scales. For the former, the following selections were made for further analysis: the +1 nucleosome, 114≤x≤168, 61.9%≤y≤80.3%, z≥0.296; the +2 nucleosome, 311≤x≤368, 67.6%≤y≤85.3%, z≥0.220; the +3 nucleosome, 506≤x≤569, 66.1%≤y≤82.8%, z≥0.174. The cutoff z values were at 1σ from the maximal z values, based upon Gaussian distribution curve fitting of xz cross sections. For the latter, the following selections were made for further analysis: 1000≤x≤4000, 80≤y≤120.

### Yeast data

*Saccharomyces cerevisiae* TSS information (Zhang and Dietrich 2005) and genome sequence were retrieved from *Saccharomyces* Genome Database (SGD, http://www.yeastgenome.org/). The paired-end MNase-Seq data (Kent et al. 2011) were download from SRA with accession SRA020615.3. Analysis was performed as with the *Tetrahymena* datasets.

### Data accession

All MNase-Seq data have been deposited at the NCBI Gene Expression Omnibus (GEO: http://www.ncbi.nlm.nih.gov/geo/) under accession GSE67394.

## Acknowledgments

WT *Tetrahymena* strain CU428 was obtained from the *Tetrahymena* Stock Center (https://tetrahymena.vet.cornell.edu/). The compiled annotations for *Tetrahymena* MAC genome sequence were obtained from the *Tetrahymena* Genome Database (<u>www.cilate.org</u>). *Tetrahymena* MIC genome sequence was obtained from the *Tetrahymena* Comparative Sequencing Project (Broad Institute of Harvard and MIT: http://www.broadinstitute.org/annotation/genome/Tetrahymena/MultiHome.html). Illumina sequencing was performed at the DNA Sequencing Core of the University of Michigan. J.X. was supported by the Natural Science Foundation of China (No. 31301930). S. G. was supported by the Natural Science Foundation of China (No. 31470064) and the Science Foundation of Shandong Province (ZR2014CQ011). S.D.T. was supported by NIH (R01 GM106024). R.E.P. was supported by grants from the Canadian Institutes of Health Research (grant number MOP13347) and the Natural Sciences and Engineering Research Council of Canada (grant number 53909). W.A. was supported by a postdoctoral fellowship from the Natural Sciences and Engineering Research Council of Canada. W.M. was supported by the Projects of International Cooperation and Exchanges, Ministry of Science and Technology of China (No. 2013DFG32390). Y.L. was supported by NSF (MCB 1411565), NIH (R01 GM087343), and the Department of Pathology at the University of Michigan.

## Supplemental Methods

### Mono-nucleosomal DNA purification by agarose gel electrophoresis

The MNase digestion was stopped by adding 10 mM EGTA, 1 mM EDTA, and 1% [v/v] SDS. After sequential RNase A and Proteinase K treatment, DNA was purified by phenol-chloroform extraction and ethanol precipitation. After agarose gel electrophoresis, mono-nucleosome sized DNA fragments (120-250 bp) were excised and purified by Zymoclean™ gel DNA recovery kit (Zymo Research).

### Mono-nucleosomal DNA purification by sucrose gradient ultracentrifugation

The MNase digestion was stopped by adding 10 mM EGTA, 1 mM EDTA, and 5 mM sodium butyrate. Solubilized chromatin was extracted by adding 0.1 % [v/v] Triton X-100 and incubating on ice for 10 min. Solubilized chromatin was separated from the nuclear pellet by centrifugation at 3,000 g for 10 min. Solubilized chromatin was fractionated by ultracentrifugation at 103,000 g for 17 hours in the sucrose gradient buffer (10 mM Tris-HCl pH 7.4, 0.1 mM EDTA, 0.1 M NaCl, 5 mM Na Butyrate, with 5%, 10%, 15%, 20%, 25% sucrose). Mono-nucleosome containing fractions were identified by OD 260/280 measurement, and validated by protein electrophoresis. DNA was subsequently purified after RNase A and Proteinase K treatment, phenol-chloroform extraction, and ethanol precipitation. 10% of the total sample was analyzed by agarose gel electrophoresis and fractions containing mono-nucleosome sized DNA were pooled.

### Illumina sequencing and data processing

Illumina sequencing libraries were prepared according to manufacturer’s instructions without the fragmentation step. Paired-end sequencing was performed using an Illumina Hiseq2000 sequencer. MNase-Seq reads of each sample were initially mapped to the genome assembly of *T. thermophila*, using TopHat2 allowing 2 edit distances including mismatch and indel (Trapnell et al. 2012). For the MAC MNase-seq data, the reads were mapped to the newest MAC genome assembly in the *Tetrahymena* genome database (TGD, (Stover et al. 2012)) (http://ciliate.org/index.php/home/downloads) ; for the MIC MNase-Seq data, the reads were mapped to the MIC genome assembly of *T. thermophile* (http://www.broadinstitute.org/annotation/genome/Tetrahymena/MultiHome.html). Unique mapped reads were extracted, and only one of any potential PCR duplicates (fragments, defined by a pair of properly mapped reads, with the same location in genome) were kept. Only mono-nucleosome sized fragments (120 to 260 bp) were analyzed. The mapped reads were visualized using Gbrowse2 (Stein 2013).

### Identification of TSS and TES in *Tetrahymena*

The published gene models (http://ciliate.org/index.php/home/downloads) do not contain information about untranslated regions (UTR), as well as TSS and TES. To obtain the relevant information, the gene models were updated using two batches of RNA-Seq data (GEO accession: GSE27971 (Xiong et al. 2012) and GSE64064). The RNA-Seq data for each sample were mapped to the MAC genome assembly using TopHat2 allowing 2 edit distances (Trapnell et al. 2012). Transcripts were independently assembled for each sample and then merged using the cufflinks program (Trapnell et al. 2012). The predicted gene models were updated using PASA (Haas et al. 2003), by aligning the assembled RNA-Seq transcripts to the genome using BLAT (Kent 2002) and GMAP (Wu and Watanabe 2005). Three rounds of PASA updating were applied, according to the suggested protocols from the PASA website (http://rice.plantbiology.msu.edu/training/Haas_PASA2.pdf). Gene models with UTR annotated in both 5’ and 3’ ends (15,841 models) were selected from the PASA updates. To avoid the overextension of the UTR, the 5’ and 3’ ends of UTR was trimmed if the coverage is less than 5× in total RNA-Seq data.

### DNA sequence-based prediction of nucleosome occupancy

Computational predictions were made using a comprehensive model built upon a dataset from nucleosomes reconstituted with yeast histones and genomic DNA in vitro (Kaplan et al. 2009). It uses dinuculeotide statistics and 5-mer statistics to compute a score for each position in the genome. These scores are used in a dynamic programming algorithm to calculate the probability for each position that it is covered by any nucleosome, which takes into account steric hindrance constraints between neighboring nucleosomes. Computational predictions were also made using the 14 DNA sequence features selected for their dominant effects on nucleosome occupancy (Tillo and Hughes 2009). Instead of the more than 2000 features used in the Segal model (Kaplan et al. 2009), this model uses only 14 features selected using a linear regression algorithm. The 14 features in this model are: GC content, slide, AAAA, propeller propeller twist, AAAT, AATA, ATAA, GAAA, ATTA, TATA, AATT, ATAT, AGAA, and AAGT. The other 16 features were created for a model based on geometrically transformed Tsallis entropy (Wu et al. 2014). Eight of the features are frequencies of all possible 3-mers using the WS alphabet (W: A/T, S:C/G). These features are calculated for both strands. The other eight features were taken from the Tsallis entropy vectors created from those features, s1 and s2.

Thirty sequence features were computed for every position in the genome using a 150 bp window sliding in jumps of 1 bp centered on the genomic position. The two structural features, slide and propeller twist, were calculated using a 75 bp sliding window instead of the 150 bp sliding window, as specified in (Tillo and Hughes 2009). Correlations for the MAC were calculated using only the scaffolds longer than 10 Kbp. Correlations for the MIC were calculated separately for IES and MDS. MDS were sequences in the MIC that aligned to sequences in the 229 MAC scaffolds that are longer than 10 Kbp. IES were sequences in the MIC genome that did not align to sequences in the MAC genome and which had no N’s in their sequence or in the 75 bp on either side of their sequence. This restriction was imposed because the computational predictions are not accurate when there are N’s in the sequence. Because of the quality of the MIC genome sequence, this excludes about half of the IES sequences.

Pearson correlations were also calculated, but we chose to report the Spearman rank correlations because, in most cases, there was a noticeable difference between the Pearson correlation and the Spearman rank correlation, indicating that, although there was a linear relationship as measured by the Pearson correlation, there was a much stronger monotonic relationship as measured by the Spearman rank correlation.

### Analysis of GC content (GC%)

GC% distributions were calculated relative to TSS or nucleosomes. For the former, the calculation was based on 15,841 well-modeled genes or specified sub-groups. For the latter, the calculation was based on either the fragment centers of the selected read pairs or the dyads of the selected called nucleosomes.

### Nucleosome distribution around splice sites

Information about splice sites was extracted from the 15,841 well-modeled genes. Only introns flanked with ≥200 bp exons on both sides were used. Distribution of called nucleosome dyads was aligned to and aggregated over all relevant splice sites. For the 5’ splice sites (GT), the position of base G is 0, and the upstream (exon) 1 kb was analyzed; For the 3’ splice sites (AG), the position of base G is 0, and the downstream (exon) 1 kb was analyzed.

## Supplemental Data

### DNA sequence features associated with gene bodies

We analyzed the nucleotide composition in gene bodies and found several prominent features (Supplemental Fig. S6A, B). First, high GC content in gene bodies, relative to inter-genic regions. With specific decrease in T% and increase in G%, the AT line and GC line were further split in gene bodies, while they were superimposed in inter-genic regions. Second, TSS anomaly, mainly manifested as dramatic oscillations of A%/T%, and to a lesser degree, G%/C%, in a narrow band 1-10 bp downstream of TSS. Third, TES anomaly, mainly manifested as dramatic oscillations of A%/T%/G%, and to a lesser degree, C%, in a narrow band 1-10 bp upstream of TES. It is worth noting that the nucleotide composition oscillations at TSS and TES strongly support their precise calling, as they would have been canceled out if TSS/TES calling in a significant fraction of the genes was offset by a single nucleotide.

The TATA box (Smale and Kadonaga 2003) was enriched around TSS (Supplemental Fig. S6C), while the termination signal (Proudfoot 2011) was enriched near TES (Supplemental Fig. S6D). The CentriMo software from the MEME suite (Bailey and Machanick 2012) was used to search for enrichment of the TATA box (TATAWAW) around the TSS and the termination signal (AATAAA) near the TES. Fifty-one percent of the well-modeled genes were found to have a TATA box around TSS (from -50 bp to 0 bp). It was enriched in the region from -31 to -14 (p<8.3e^-40^). Fifty percent of the well-modeled genes were found to have a termination signal (AATAAA) around TES (from -50 to 50). This signal was enriched ending at the TES (p<3.3e^-10^).

### Nucleosome positioning in the rDNA mini-chromosome in *MAC*

In addition to those associated with transcription, there were other trans-determinants affecting nucleosome positioning in MAC. We focused on the rDNA mini-chromosome, which is processed from a single copy in MIC by excision, palindrome formation, and de novo telomere addition, and highly amplified in MAC (Supplemental Fig. S10A). Periodic distribution of MNase-digested DNA fragments were found around the rDNA replication origins (Supplemental Fig. S10B). Interestingly, this region was only recovered from samples with light MNase digestion. Comparison between the samples prepared by gel purification and sucrose gradient ultracentrifugation showed that the footprints in the two rDNA replication origins (ori1 and ori2) were prominent only in the former, but not the latter, strongly supporting their non-nucleosomal nature and consistent with binding of the origin recognition complex (ORC) and other key factors for replication initiation (Kapler 1993). With the exception of a nucleosome position at the center of the palindromic rDNA, we recovered all positioned nucleosomes (Supplemental Fig. S10B), as reported in classical MNase digestion analysis of 5’ non-transcribed sequence (5’ NTS) of rDNA (Palen and Cech 1984). We also recovered a positioned nucleosome in the rDNA sub-telomeric region ∽150 bp away from the G4T2 telomere repeats (Supplemental Fig. S10C), consistent with classical MNase digestion analysis of 3’ NTS and the sub-telomeric region (Budarf and Blackburn 1986). Interestingly, the degrees of translational positioning of these nucleosomes were significantly lower than many in actively transcribed gene bodies, suggesting that transcription has by far the strongest impact on the chromatin environment.

### Supplemental figure legends

**Supplemental Figure S1.**
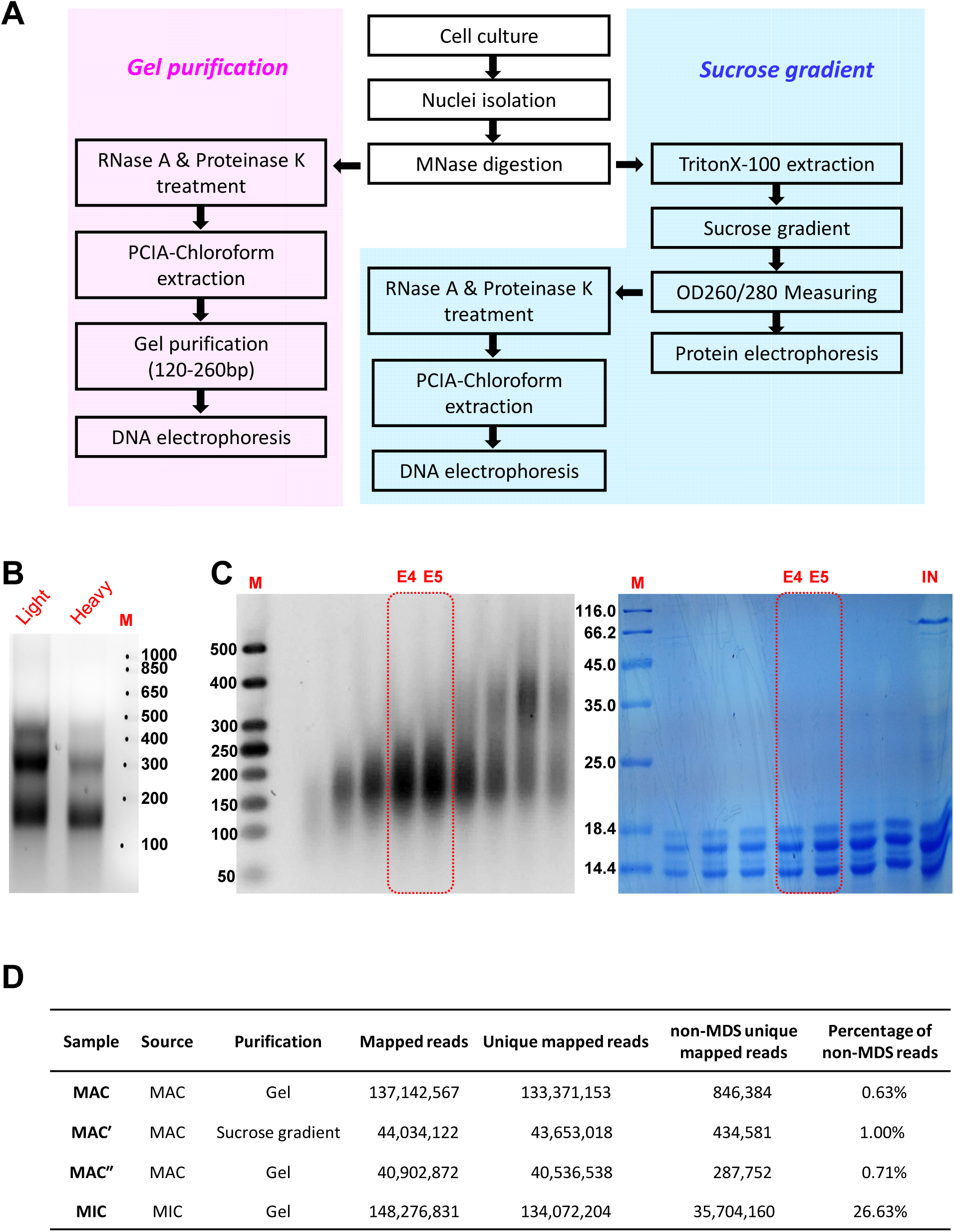
Mapping nucleosome distribution in *Tetrahymena* MAC and MIC by MNase-Seq, overview. A) Sample preparations for MNase-Seq. After MNase digestion, mono-nucleosome sized DNA fragments are purified by either agarose gel electrophoresis or sucrose gradient ultracentrifugation. B) Nucleosome ladders after light or heavy MNase digestion of *Tetrahymena* MAC. Note the ∽200 bp NRL. M: markers for DNA fragment length, as indicated (bp). C) Fractionation of the MNase solubilized chromatin by sucrose gradient ultracentrifugation. Left, DNA analysis of fractions by agarose gel electrophoresis. M: markers for DNA fragment length, as indicated (bp). E4/E5: the fractions enriched for the mono-nucleosome, used for MNase-Seq (sample MAC′, see below). Right, protein analysis of fractions by SDS-polyacrylamide gel electrophoresis. M: markers for protein molecular weight, as indicated (kDa). IN: 2% input. E4/E5: the fractions enriched for the mono-nucleosome, demonstrated by the predominant staining of histones. D) Basic characteristics of the MNase-Seq samples for *Tetrahymena* MAC and MIC.

**Supplemental Figure S2.**
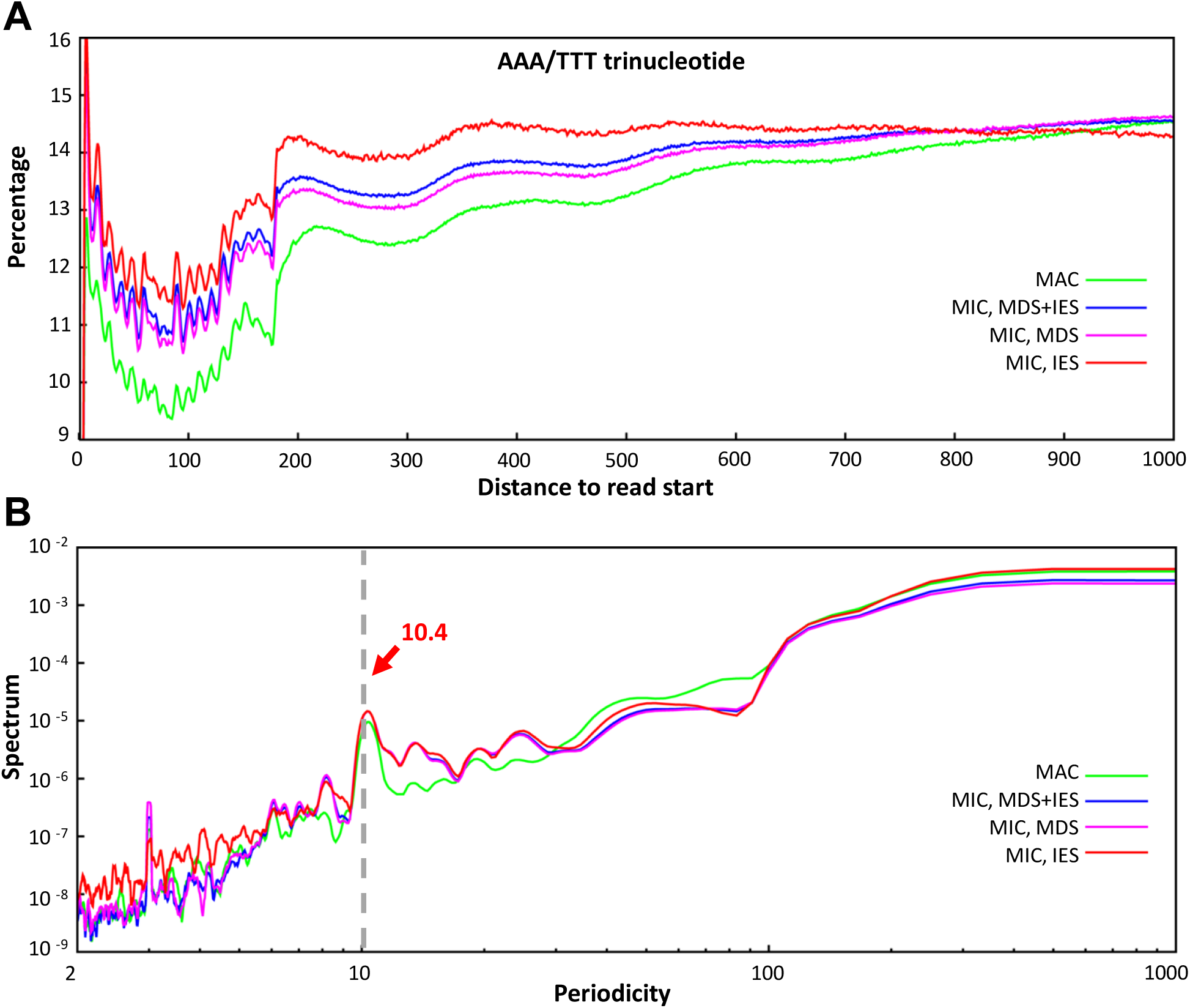
Conservation of rotational nucleosome positioning in *Tetrahymena* MAC and MIC. A) Periodic AAA/TTT trinucleotide associated with the nucleosome. Composite distribution of AAA/TTT trinucleotide frequency is aligned to the start position of all MNase-Seq reads and calculated for 1 kb downstream. B) 10.4 bp periodicity of nucleosome-associated AAA/TTT trinucleotide, revealed by the spectrum analysis. The AAA/TTT trinucleotide frequency data above are subjected to the spectrum analysis, also known as discrete Fourier transform, using “spectrum” function in R. A spectrum peak (gray dashed line) indicating a 10.4bp periodicity is detected with consistent position and height in the MAC and MIC datasets.

**Supplemental Figure S3.**
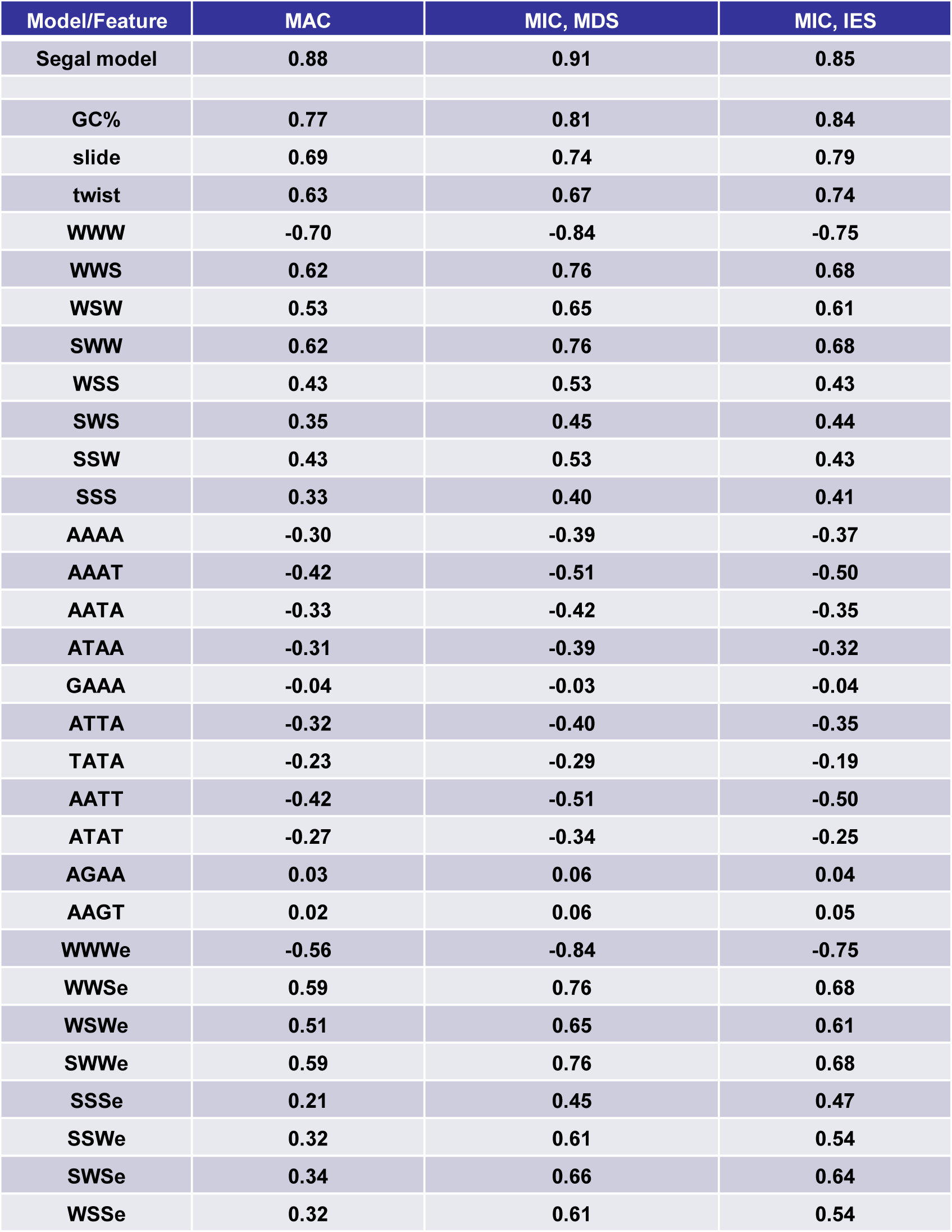
Spearman’s rank correlations between nucleosome occupancy levels calculated from the MNase-Seq datasets and predicted by computational modeling. The Segal model is the comprehensive computational model (Kaplan et al. 2009). GC%, slide, propeller twist and the eleven 4-mer features are from the 14 feature model (Tillo and Hughes 2009). The eight 3-mer features (W: A/T, S: C/G) as well as the eight Tsallis entropy features are as described (Wu et al. 2014). See Methods and Supplemental Methods for details.

**Supplemental Figure S4.**
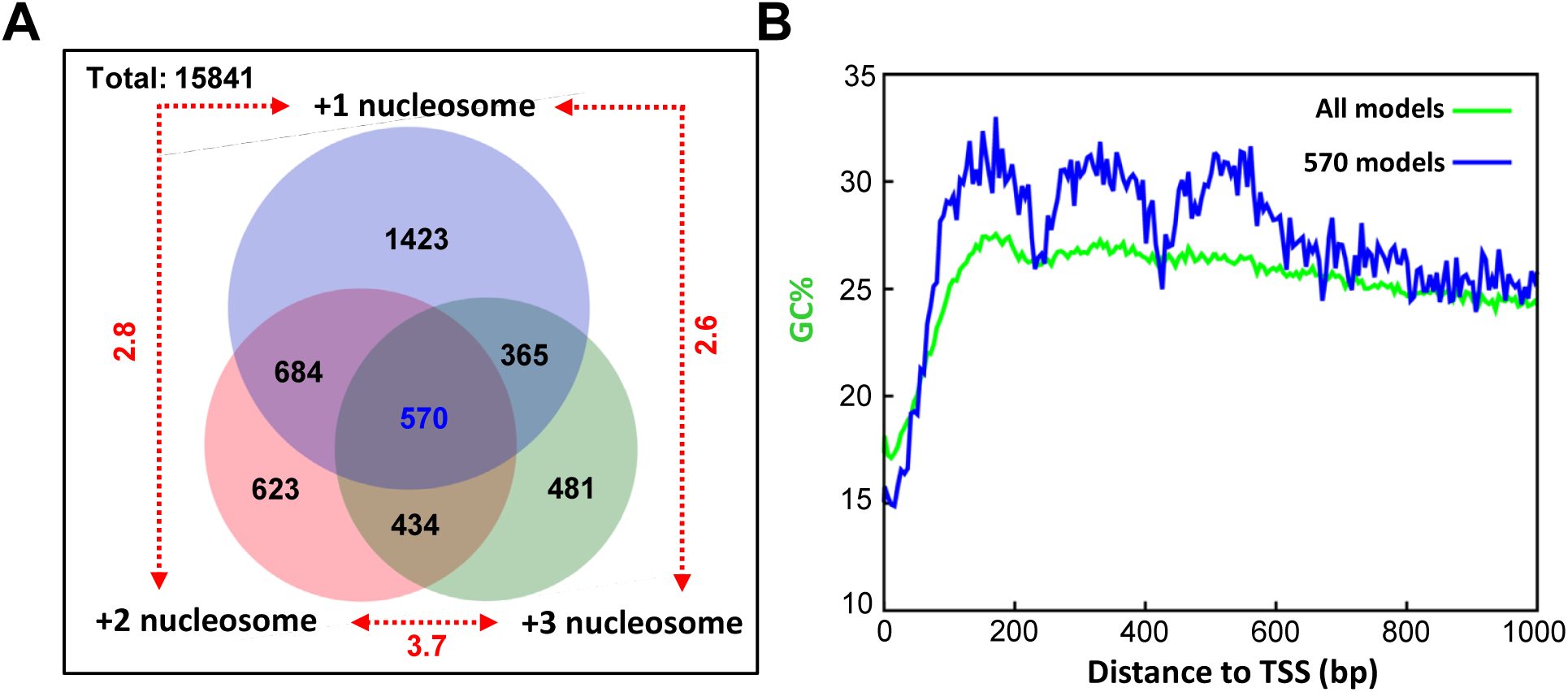
Comparison of GC content distribution relative to TSS in different gene sets. A) Overlap of genes with the well-positioned +1, +2, or +3 nucleosomes. The logical relationships between the three sets containing the well-positioned +1, +2, or +3 nucleosomes (as in Figure 3A) are illustrated by Venn diagram. The numbers of genes in each region of the Venn diagram are indicated. High representation factors (red, next to dashed lines) for each intersection indicate strong tendencies for the well-positioned +1, +2, or +3 nucleosomes to cluster, i.e., to be found in the same gene (p«1e-200 for all). The intersection between all three sets is further analyzed (see below) B) GC% distribution relative to TSS, comparing all 15,841 well-modeled genes to 651 genes with well-positioned +1, +2, and +3 nucleosomes (see above). Note the prominent GC% peaks downstream of TSS in the latter but not the former.

**Supplemental Figure S5.**
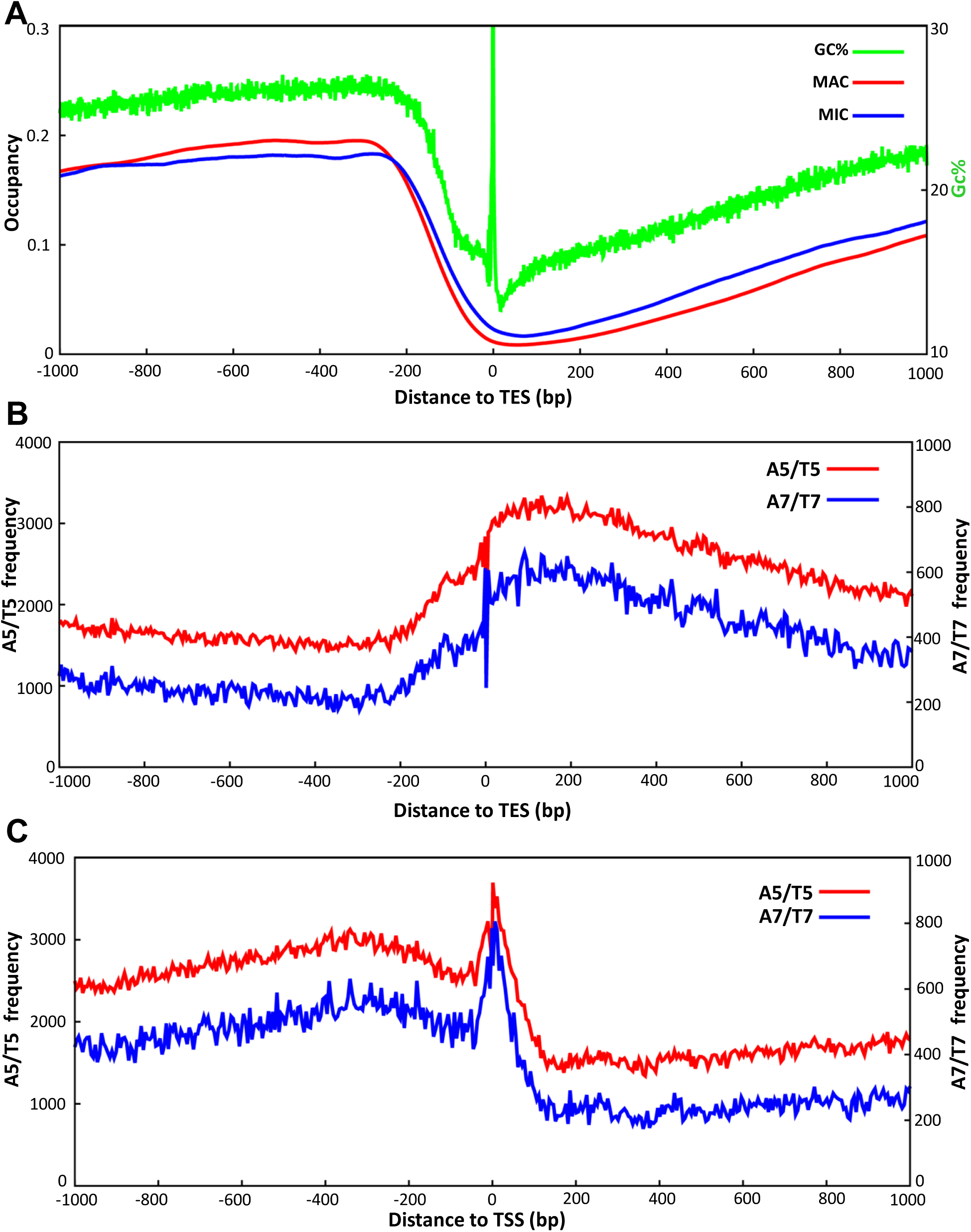
Additional analyses of TSS and TES. A) Nucleosome-depleted regions (NDR) around TES. Distributions of MNase-Seq fragment centers are calculated around TES (±1 kb) using the MAC (red) and MIC (blue) samples, with frequency normalized to per million sequenced fragments. GC% distribution (green), also aligned to TES, is superimposed. B) Distributions of poly (dA:dT) tracts relative to TES. A5/T5 and A7/T7 distributions are calculated around TES (±1kb) using the 5bp bin. C) Distributions of poly (dA:dT) tracts relative to TSS. A5/T5 and A7/T7 distributions are calculated around TSS (±1kb) using the 5bp bin.

**Supplemental Figure S6.**
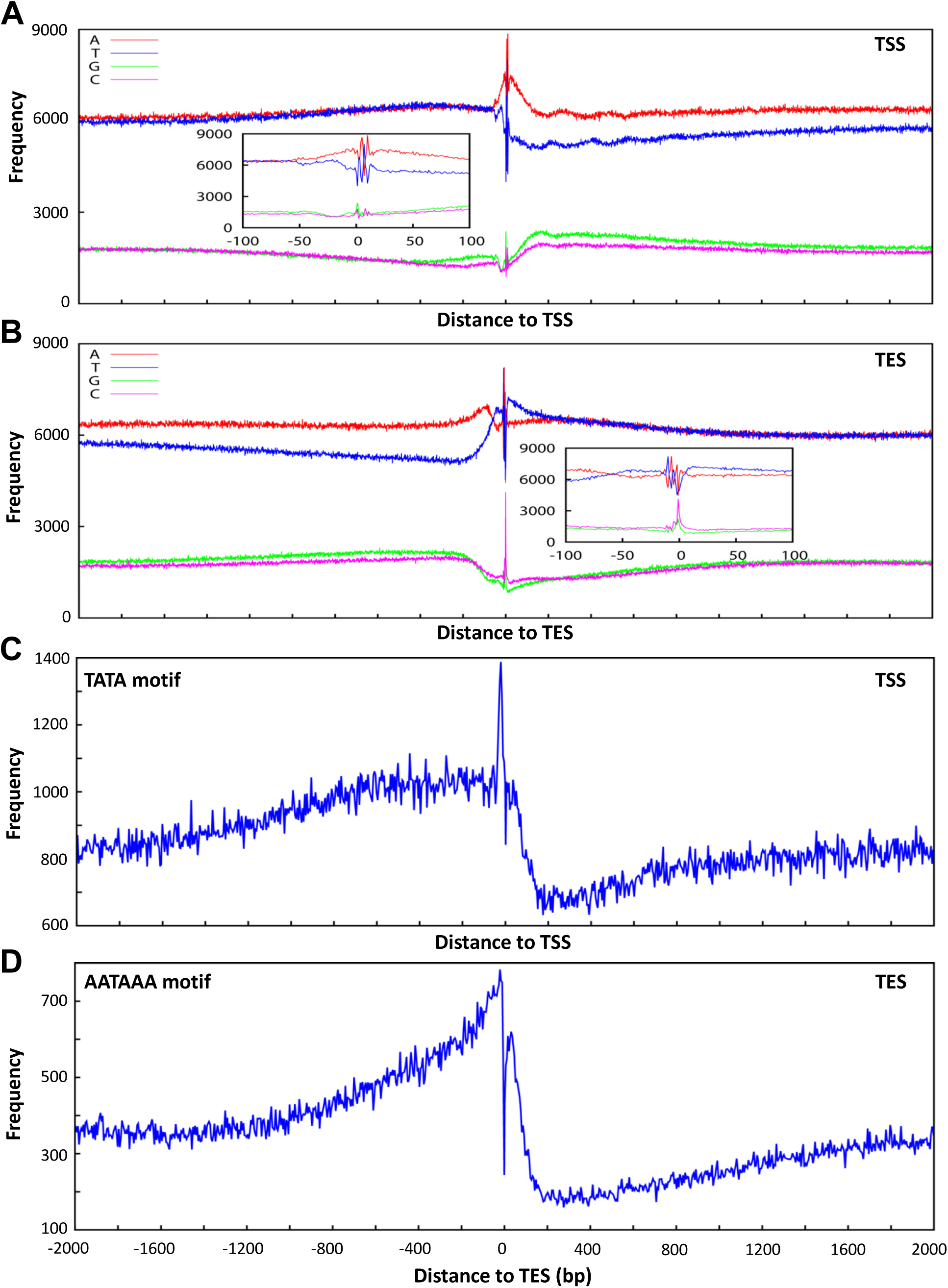
Base composition anomalies in regions around TSS and TES. See Supplemental Data for more details. A) Base composition around TSS (±2 kb), calculated from 15,841 well-modeled genes. Inset: zoom in on the ±100 bp region. B) Base composition around TES (±2 kb), calculated from 15,841 well-modeled genes. Inset: zoom in on the ±100 bp region. C) The distribution of TATA, potentially corresponding to the TATA box (Smale and Kadonaga 2003), around TSS (±2 kb). See Supplemental Data for more details. D) The distribution of AATAAA, potentially corresponding to the polyadenylation signal (Proudfoot 2011), around TES (±2 kb). See Supplemental Data for more details.

**Supplemental Figure S7.**
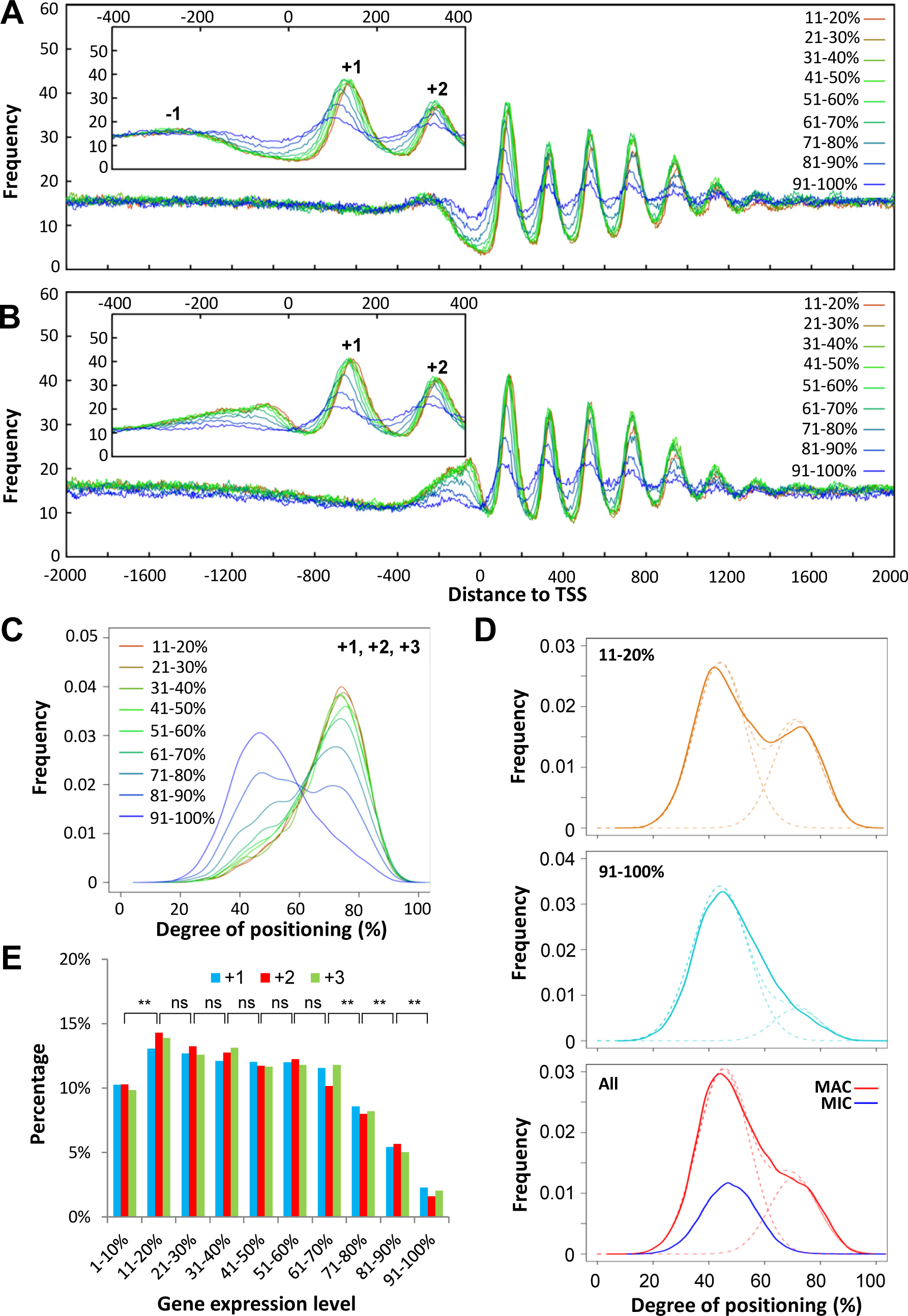
Nucleosome arrays in gene bodies are affected by transcription levels. A) Stereotypical nucleosome arrays in genes with different transcription levels. The MNase-Seq sample (MAC′) was generated by light MNase digestion and purified by sucrose gradient ultracentrifugation. 15,841 well-modeled genes are ranked from high to low by their expression levels and divided into 10 quantiles. Distribution of MAC MNase-Seq fragment centers around TSS is aggregated over all genes in a quantile. Normalized distributions for quantile 2-10 are plotted. Inset: zoom in around TSS (-400 to 400 bp). Note that the amplitudes of the periodic nucleosome distribution in gene bodies decrease with decreasing expression levels. Additionally, NDR between the +1 and -1 nucleosomes are increasingly occupied, likely with labile nucleosomes, with decreasing gene expression. B) Stereotypical nucleosome arrays in genes with different transcription levels. The MNase-Seq sample (MAC″) was generated by light MNase digestion and gel purified. Note that the non-nucleosomal binding events upstream of TSS—attributable to specific transcription factors, the general transcription machinery, as well as transcription-associated ATP-dependent chromatin remodelers—decrease with decreasing expression levels. C) Translational nucleosome positioning in genes with different transcription levels. Nucleosome distribution is aggregated over all called +1, +2, and +3 nucleosomes of a specified quantile. Normalized distributions for quantile 2-10 are plotted. Note that as transcription levels decrease, degrees of translational positioning of associated nucleosomes also decrease, with reduction in the well-positioned nucleosome peak and increase in the delocalized nucleosome peak. D) Nucleosome delocalization in genes with little or no expression. The nucleosome distribution curve (solid line)—plotted according to their degrees of translational positioning—can be decomposed into two peaks of Gaussian distribution (dashed line), with the left peak representing more delocalized nucleosomes and the right peak well-positioned nucleosomes. Top panel, quantile 2 contains highly expressed genes, associated with many well-positioned nucleosomes; middle panel, quantile 10 contains genes with little or no expression, associated with mostly delocalized nucleosomes; bottom panel, nucleosome distributions for all called nucleosomes in MAC, and MIC nucleosomes corresponding to the well-positioned nucleosomes in MAC (as in Fig. 1D, left panel). Note the correspondence between the left and right peaks in all panels. See Supplemental Data for more details. E) Well-positioned nucleosomes are significantly reduced in genes with very high or very low expression levels. The +1, +2, and +3 nucleosomes (as in Fig. 3A) are separated into 10 quantiles ranked from high to low by expression levels of associated genes, shown in percentage. The p values for significantly different percentages between neighboring quantiles (denoted by ^**^) are 0.00077 (between quantile 1 and 2), 0.00552 (7 and 8), 0.00032 (8 and 9), and 0.00026 (9 and 10). The others are non-significant (ns; p>0.05).

**Supplemental Figure S8.**
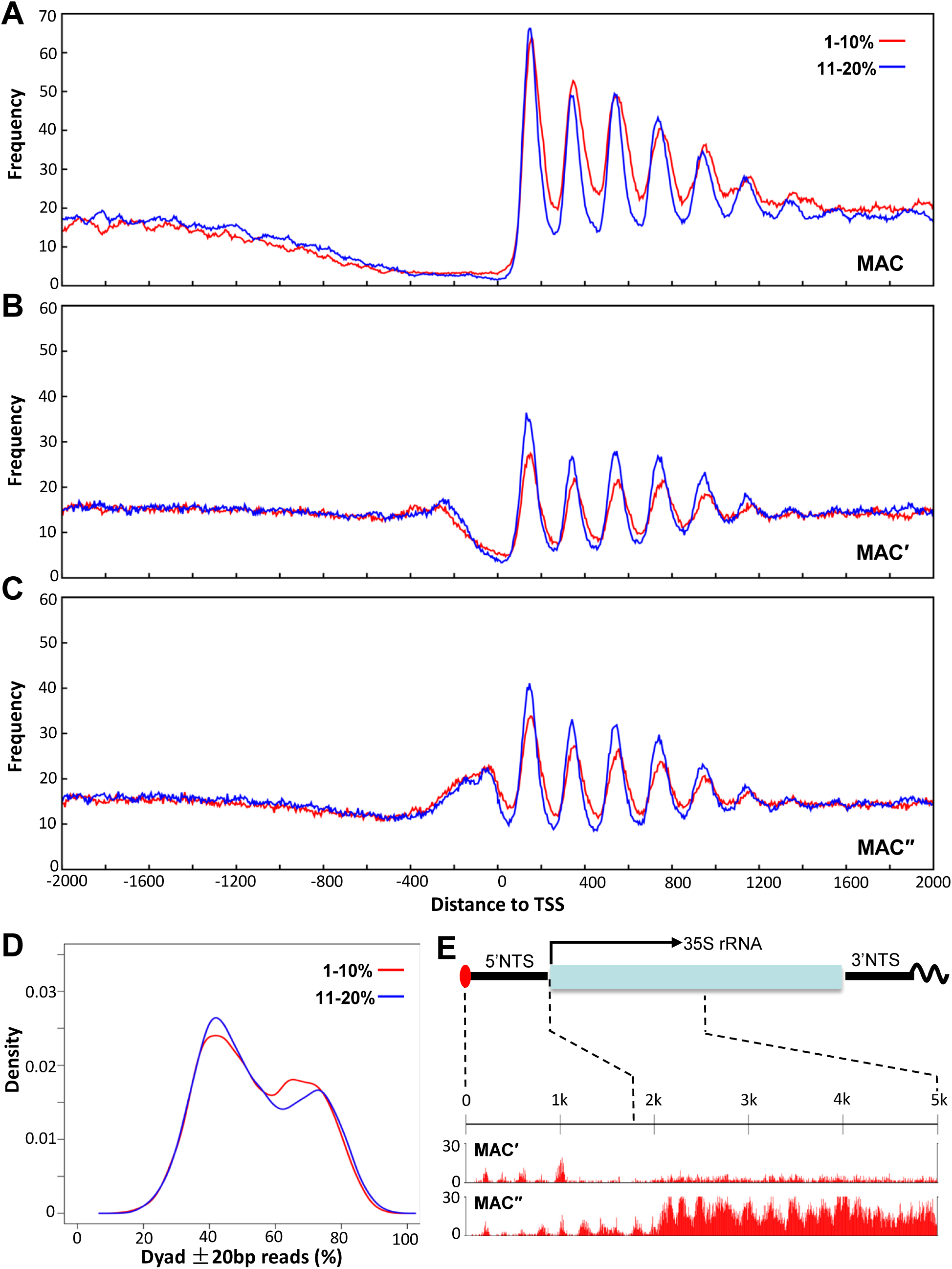
Disrupted nucleosome arrays in genes with the highest expression levels. A-C) Comparison of stereotypical nucleosome arrays in genes from the top 2 quantiles ranked by their expression levels (defined in Fig. 5A). MNase-Seq results from the MAC (A), MAC′ (B), and MAC″ (C) are aligned to TSS, and normalized to per million reads per gene. Note the reduced amplitudes of the periodic nucleosome distribution in the highest expression quantile compared with the second quantile. D) Comparison of translational positioning of nucleosomes in genes from the top 2 quantiles ranked by their expression levels (defined in Fig. 5A). Nucleosome distribution is aggregated over all called nucleosomes in gene bodies of a specified quantile (defined in Supplemental Fig. S7C). Normalized distributions for quantile 1 and 2 are plotted. Note that degrees of translational positioning in quantile 1 are lower than quantile 2, with reduction in the well-positioned nucleosome peak and increase in the delocalized nucleosome peak. F) Depletion of positioned nucleosomes in the transcribed region of the rDNA mini-chromosome. Note the positioned nucleosomes in the 5’ non-transcribed sequence (5’ NTS), which are detectable in the MAC′ and MAC″ samples with light MNase digestion, but not the MAC sample with heavy MNase digestion (not shown).

**Supplemental Figure S9.**
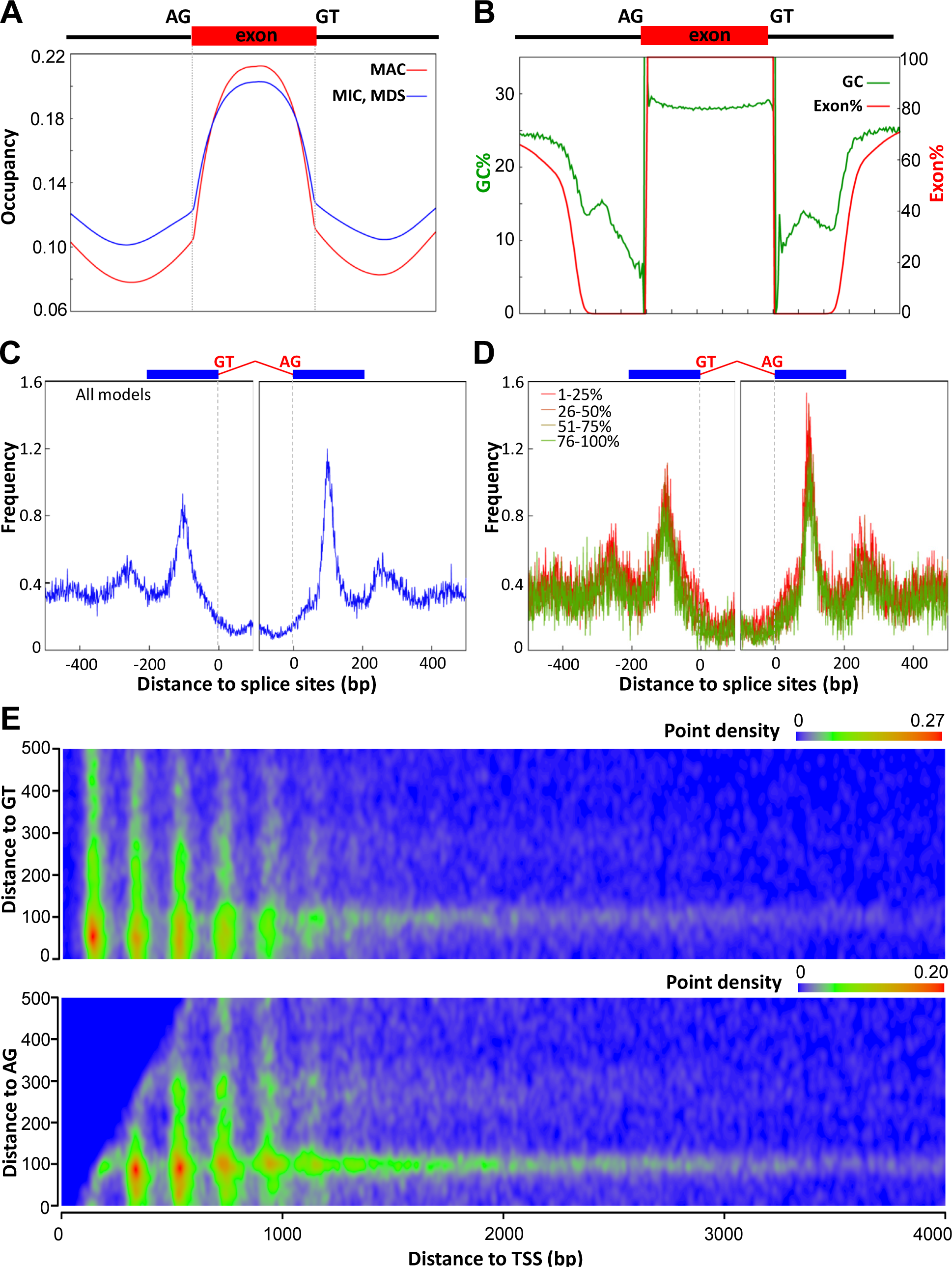
Additional analyses of the nucleosome distribution near splice sites. A) Nucleosome occupancy in and around internal exons. Internal exons (≥100 bp) from 15,841 well-modeled genes are divided into 100 parts, and then extended by100 bp in both directions. The average occupancy at each part in the exons, as well as the occupancy at each bp in the flanking regions is calculated from the MAC (red) and MIC (blue) MNase-Seq results, aggregated, and normalized to per internal exon. B) GC% and exon% in and around internal exons. The distribution of exon% (red)—calculated as the probability that a particular position containing exons—as well as GC% (green) are plotted in the frame of reference described above. C) Nucleosome distribution relative to splice sites. Introns flanked with exons at least 200 bp in length on both sides are analyzed. Distribution of called nucleosomes (dyad) is aligned to and aggregated over all qualified splice sites (5’: GT, 3’: AG). D) Expression levels have no conspicuous effect on nucleosome distribution relative to splice sites. The intron-containing genes are ranked from high to low by their expression levels and divided into quartiles. Distribution of called nucleosomes (dyad) is aligned to and aggregated over all qualified splice sites (5’: GT, 3’: AG) in a specified quartile, and normalized to per splice site. E) Distribution of called nucleosomes with high degrees of translational positioning (≥0.5). The plots are otherwise the same as Fig. 6A. Note that the grid-like pattern defined by nucleosomes positioned relative to TSS and splice sites is preserved.

**Supplemental Figure S10.**
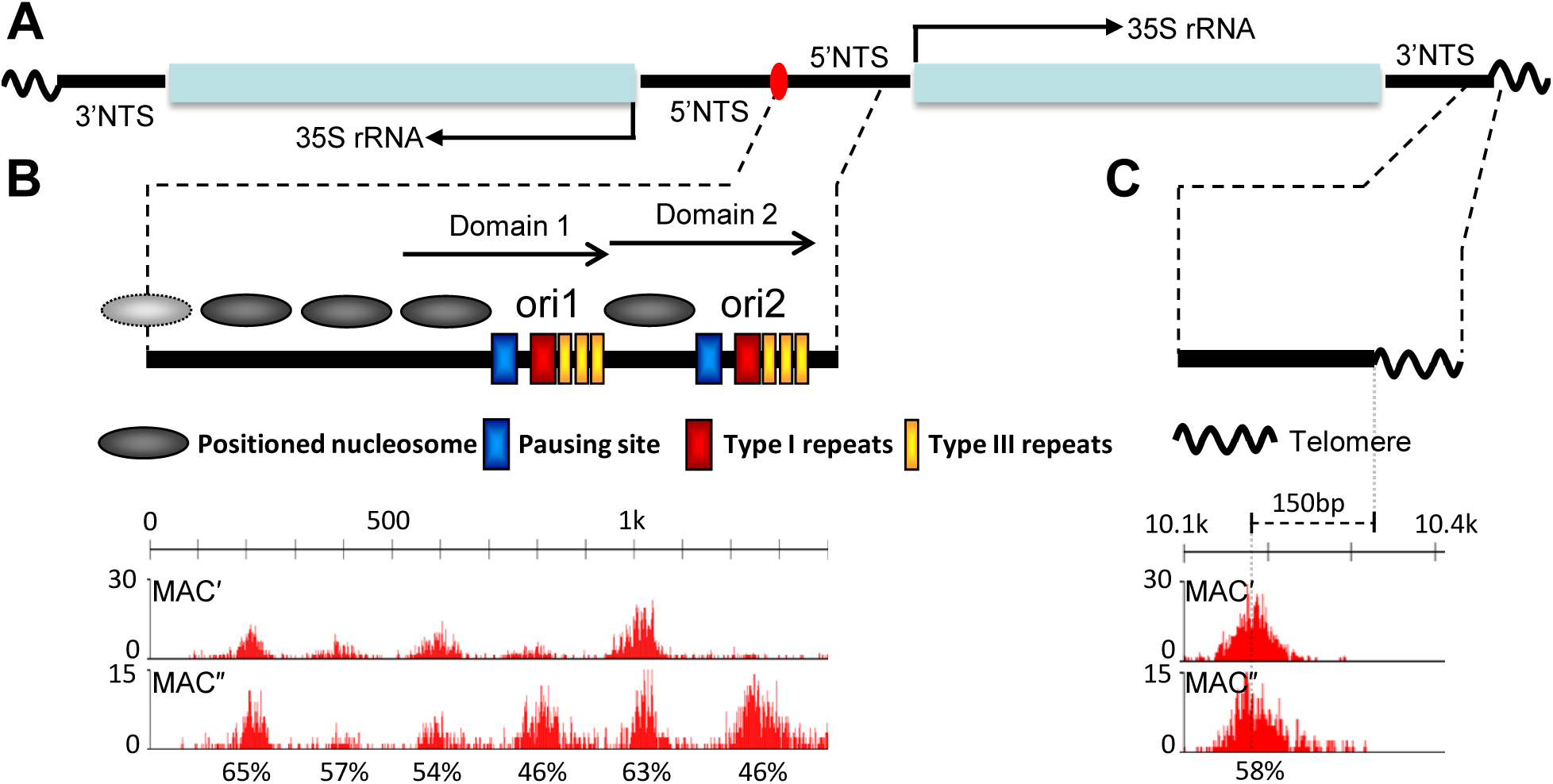
Nucleosome distribution in the rDNA mini-chromosome. See Supplemental Data for more details. A) Schematic for the palindromic rDNA mini-chromosome in *Tetrahymena* MAC. B) Positioned nucleosomes in the rDNA 5’ non-transcribed sequence (NTS), as established by the classical MNase mapping (Palen and Cech 1984). The nucleosome positioned at the rDNA center is not detected, probably due to sequencing difficulties with long palindromes. The other positioned nucleosomes are only detected in samples with light MNase digestion, MAC′ and MAC″. The degrees of translational positioning are calculated for MAC″, as indicated under the corresponding peaks of fragment centers. See Supplemental Methods for details. C) A positioned nucleosome in the rDNA 3’ NTS, as previously demonstrated (Budarf and Blackburn 1986). This positioned nucleosome is only detected in samples with light MNase digestion, MAC′ and MAC″. The degree of translational positioning for this sub-telomeric nucleosome is calculated for MAC″, as indicated under the corresponding peak of fragment centers. See Supplemental Methods for details.

### Supplemental files

Supplemental File S1. A list of called nucleosomes in *Tetrahymena* MAC.

Supplemental File S2. A list of well-modeled genes in *Tetrahymena*.

Supplemental File S3. A list of IES in *Tetrahymena*.

Supplemental File S4. A list of MDS in *Tetrahymena*.

